# Evolution of *Wolbachia* male-killing mechanism within a host species

**DOI:** 10.1101/2025.01.13.632718

**Authors:** Hiroshi Arai, Arman Wijonarko, Susumu Katsuma, Hideshi Naka, Daisuke Kageyama, Emily A. Hornett, Gregory D. D. Hurst

## Abstract

Male-killing bacterial symbionts, prevalent in arthropods, skew population sex ratios by selectively killing male progeny, profoundly impacting ecology and evolution of their hosts. Male-killing is a convergently evolved trait, with microbes evolving diverse male-killing mechanisms across host species with widely divergent sex determination pathways. A common evolutionary response to MK presence is the spread of suppressor mutations that restore male survival. In this study, we demonstrate evolution of a novel male-killing mechanism that is insensitive to an existing male-killing suppressor. *Hypolimnas bolina* butterflies from Yogyakarta, Indonesia, showed extreme female biased population sex ratio associated with high prevalence of a male-killing *Wolbachia*. This strain, *w*Bol1Y, shared a very recent common ancestor with the previously characterized ‘suppressed’ male-killing strain in the species, *w*Bol1, but retained its male-killing ability in the presence of the male-killing suppressor. The genome of *w*Bol1Y differed from the suppressed *w*Bol1 in carrying an additional prophage that included strong candidate genes for male-killing. *In vitro* and *in vivo* data demonstrated that *w*Bol1Y feminized splicing and expression of lepidopteran sex determination pathway genes, and that the gene *Hb-oscar* – present on *w*Bol1Y’s unique prophage insert – was sufficient to disrupt the male sex determination pathway. Our study demonstrates the diversity of male-killing mechanisms is a product both of interaction with varying insect sex determination systems and evolution of male-killing within a host species. Our data indicate male-killer and host may be involved in escalating arms races, where spread male-killing suppression drives evolution of additional systems that reestablish male killing by the symbiont.

## Introduction

Heritable microorganisms are common in arthropods, and have diverse ecological and evolutionary impacts on their host (1, 2). In many cases they are beneficial partners, providing a range of services, for instance extending anabolic capacity (3), providing defence against natural enemies (4, 5), and improving desiccation resistance and thermal tolerance (6). These services impact host ecology (for instance, defining the niche of the species), and host evolution, as innovations fueling radiation (7). Contrastingly, the strict maternal inheritance of these microbes has in many cases driven the evolution of sex ratio distorting ‘manipulations’, where the symbiont enhances the production of female progeny of their host through which they can transmit, at the expense of males (2). Principle of these is the phenotype of male killing (MK), where diverse microbes have evolved to kill male hosts that inherit them, most commonly during embryogenesis (8, 9). Male-killers can have profound ecological and evolutionary impacts, through altering the population sex ratio, and through selection on the host to restore male survival. Indeed, the intense nature of Fisherian selection makes the spread of suppressors of MK activity some of the most rapid events of natural selection that have been observed in natural populations (10–15).

MK symbionts are observed in arthropod hosts with a wide diversity of sex determination systems, both from a karyotypic and functional genetic perspective (2, 9). To achieve MK, bacterial endosymbionts commonly directly ‘manipulate’ the host’s sex determination systems (16–22). For example, some MK *Spiroplasma* and *Wolbachia* endosymbionts impair or utilize the dosage compensation system, which regulates the expression levels of sex chromosome-linked genes (17–19, 21). MK *Wolbachia* can also ‘feminize’ the male sex determination cascades, as observed in some species of moth (17, 22). In the haplodiploid wasp, *Nasonia vitripennis*, the bacterium *Arsenophonus* inhibits the formation of maternal centrosome (23). Recent functional analyses indicate that the causative factors employed by male-killers are also diverse (16, 19, 20, 24, 25). This functional diversity can be considered a case of convergent evolution, with a similar MK phenotype being established in varying sex determining contexts through different microbial genes and systems.

A further possibility is that the mechanism of MK additionally diversifies within a species. MK symbionts drive host evolution as antagonists to be avoided because MK imposes a fitness cost on their host (26). This selection pressure has led to the spread of suppressor mutations that restore male survival (10–15). As a case of antagonistic coevolution (27), selection is then expected to act on the symbiont to escape suppression. Changes leading to suppression insensitivity would likely represent either enhancement of the existing MK system or addition of new systems, providing a pathway in which the diversity of MK mechanism is driven intra-specifically, as part of an arms race.

In this study, we present evidence that MK mechanisms have evolved within a host species that has a circulating suppressor of MK. Our focal system is the nymphalid butterfly *Hypolimnas bolina*, that has undergone drastic changes in its sex ratio in the last 100 years. The species carries an MK *Wolbachia* strain, *w*Bol1, that killed male embryos late in development (28, 29). In Samoa and Fiji, *H. bolina* exhibited a marked female-biased population sex ratio (100:1) in the 1920s (30, 31) that was retained in Samoa until 2000 (28, 29, 32). However, the sex ratio suddenly recovered in 2005-2006 associated with the rapid spread of suppressors against *Wolbachia*-induced MK (10, 12). In Southeast and East Asia, the sex ratio of the butterfly was also skewed towards females until the 1980s (33, 34), and the skewed sex ratio was observed to recover to 1:1 in many Asian populations around the 1980s-1990s (35, 36). These observations evidence the spread of a suppressor that has silenced the MK *Wolbachia w*Bol1 over a very broad geographical area, allowing infected males to survive.

Our study was motivated by an initial observation of persistent extreme female-biased population sex ratios in a population of *H. bolina* butterflies in Yogyakarta (Jawa, Indonesia), that contrasts with the widespread host suppression of MK in Asian *H. bolina* populations, including that observed on the adjacent island of Borneo (37, 38). We provide evidence that MK in Yogyakarta was induced by a variant of *Wolbachia w*Bol1 that was insensitive to the MK suppressor widely established in *H. bolina*. Further, we establish that this ‘non-suppressible’ strain carried an additional MK factor, *Hb-oscar*, through acquisition of a prophage element. We conclude that the mechanism of MK evolves not only in the context of the diverse sex determination systems found across host species, but also evolves within a host species.

## Results

### Extreme prevalence of MK in a Yogyakarta population of *H. bolina*

We observed a significantly female biased population sex ratio in *H. bolina* butterflies collected in Yogyakarta over 8 years since 2016 (field collections: 209 females vs. 4 males in total, P = 0.000, binomial test). This strong female bias was redolent of the historically observed strongly skewed sex ratio in Samoa (31) and Fiji (30) (**Fig. 1A**). In contrast, field-collected Japanese *H. bolina* showed a male-biased population sex ratio from 2021 to 2023 (16 females vs. 162 males in total, P = 0.000). Mitsuhashi et al. (36) and Ikeda (35) suggested that male-biased sex ratios observed in field-collected samples reflects the territorial behavior of males and the relatively low activity of females. In contrast, in the female-biased Yogyakarta population, females were notably active, displaying territorial behavior by chasing other individuals, while males were mostly hidden in the bushes, with very little observed flight activity. This sex role reversal echoed that observed in Samoa, and in *Acraea encedon* with high prevalence MK *Wolbachia* in Uganda (29, 39).

**Fig. 1.**
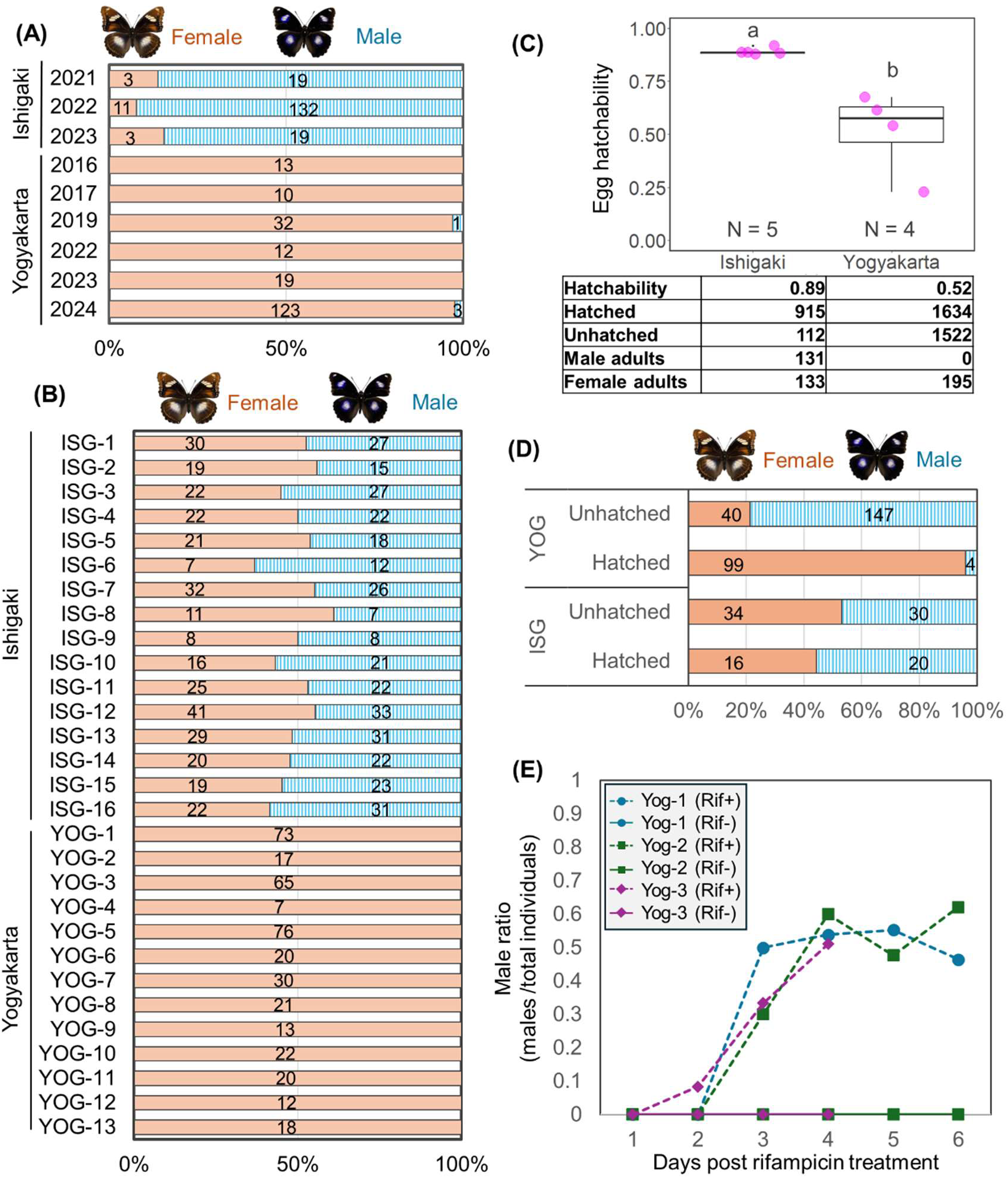
**Characterisation of the MK traits in *H. bolina*** (A) Sex ratio of field-collected *H. bolina* butterflies in Ishigaki, Japan and Yogyakarta, Indonesia. Orange: female; blue: male. (B) Sex ratio of offspring produced by field-collected females. Orange: female; blue: male. (C) Hatchability of embryos in all-female brood (Yogyakarta) and brood with normal sex ratio (Ishigaki). (D) Sex ratio of unhatched embryos and hatched 1st instar larvae. Unhatched embryos were dissected 8 days after oviposition. (E) The changes in sex ratio of the offspring of mothers treated with the rifampicin-supplemented diet. The x-axis indicates days after the start of rifampicin treatment, and the y-axis the sex ratio of the offspring laid on that date. Females from three all-female lines were treated with the same procedure. Note, female YOG-3 (Rif+) died on day 4, no observation was made in the days 5 and 6 for YOG-3 (Rif-).

The capacity of females to acquire mates, and the sex ratio produced by females, also varied between the two populations. Field-collected Japanese females (n = 16) were already mated and oviposited fertilised eggs, and the offspring showed a normal sex ratio (344 females and 345 males in total, P = 0.500, binominal test) (**Fig. 1B**). In contrast, more than a half of the field-collected females in Yogyakarta were unmated (24 out of 37 females), and females that oviposited fertilised eggs (n = 13) produced only female adult progenies (394 females and 0 males in total, ranging from 7-73 per female) (**Fig. 1B**). The egg hatchability of all-female broods in Yogyakarta (51.3 ± 1.71%, n = 4) was significantly lower than that of the normal sex ratio Japanese lines (89.0 ± 1.55%, n = 5) (Wilcoxon test, P = 0.015) (**Fig. 1C**).

To confirm the sex ratio of the hatched and unhatched individuals in the all-female broods (from Yogyakarta), we determined the sex of dead embryos and hatching individuals by observing the presence/absence female-specific heterochromatin W chromosome (presence: female; absence: male). While unhatched embryos were male-biased (40 females vs. 147 males, male ratio: 78.6%; P = 0.000, binomial test), hatched neonates were female-biased (99 females vs. 4 males, male ratio: 3.88%; P = 0.000) (**Fig.1D**). Furthermore, rifampicin treatment of female butterflies from all-female lines restored adult male survival in their offspring (**Fig.1E**). These results are consistent with the role of endosymbiotic bacteria in causing the MK phenotype.

### Closely related *Wolbachia* strains in *H. bolina* exhibited phage-linked genomic differences

All *H. bolina* lines established above were infected with MK *w*Bol1 *Wolbachia*, notwithstanding the host sex ratio produced. The *w*Bol1 *Wolbachia* exhibited identical typing gene sequences to those of the previously described *w*Bol1, with MLST ID 40 of Charlat et al. (40) and MLST ID 270 of Mitsuhashi et al. (32), collected from French Polynesia and Japan, respectively (**Fig. 2A**). 16S rRNA metagenomic sequencing revealed that the *w*Bol1 *Wolbachia* dominated in both lines (**Table S1**). Other than *Wolbachia*, no known MK bacteria specific to the all-female line were identified (**Table S1**).

**Fig. 2.**
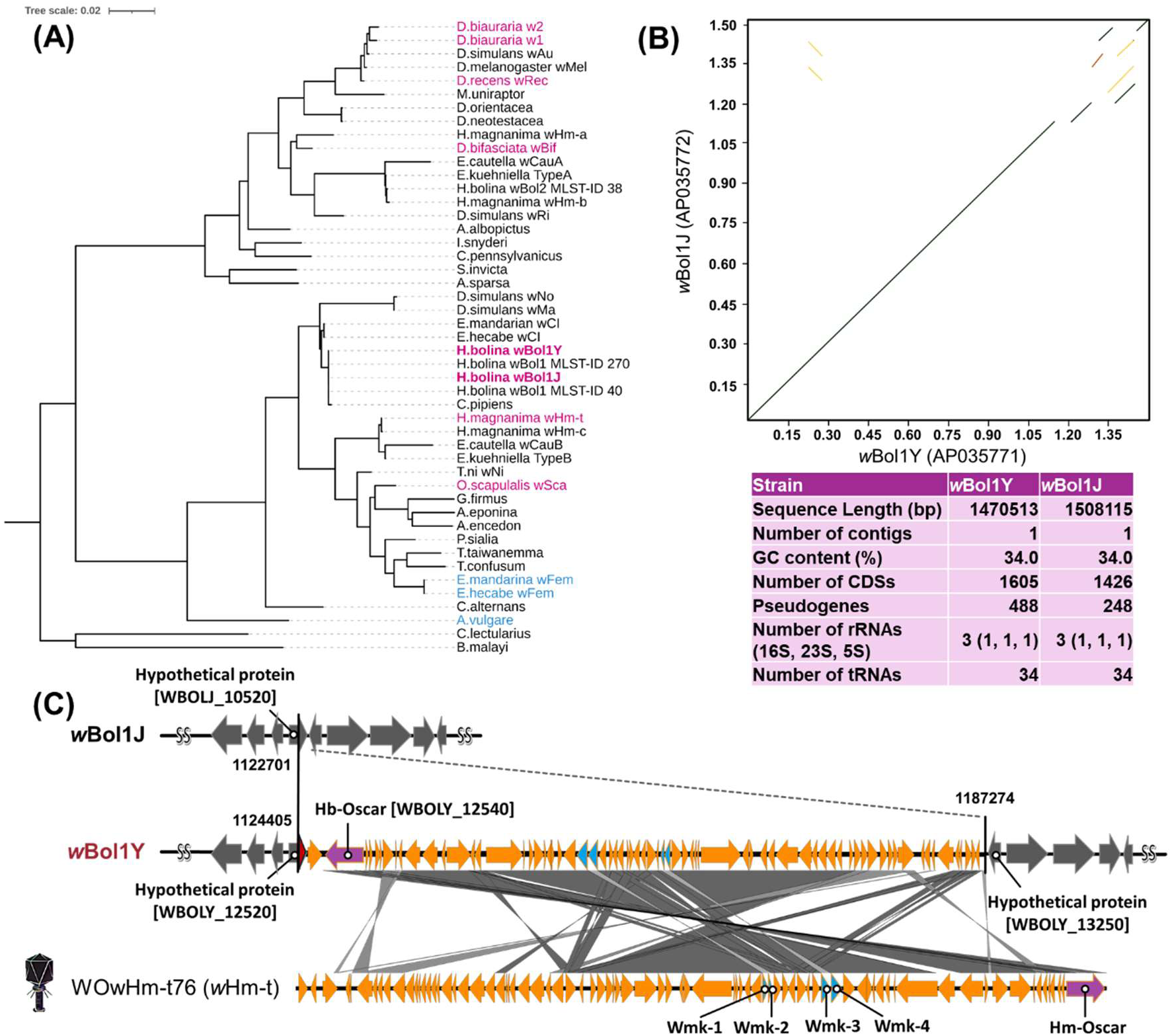
**Comparative genomics of *w*Bol1 *Wolbachia*** (A) Phylogenetic tree based on the *Wolbachia* typing genes *wsp* and MLST. MK and feminising *Wolbachia* strains are highlighted in magenta and blue, respectively. (B) Harr plot of the genomes of *w*Bol1J (currently non-MK, Ishigaki) and *w*Bol1Y (MK, Yogyakarta). Sequence similarity of genomes is highlighted in green (>80% identity), yellow (50-80% identity), and orange (30-50% identity). General information about the two strains is given below the Harr plot. Pseudogenes indicated are coding sequences that showed substitutions or frameshifts compared to the marker *Wolbachia* gene sets available from DFAST. (C) Prophage insertion into *w*Bol1Y. A *w*Bol1Y-specific prophage insertion was identified in the *w*Bol1Y genome (1124-1187 kb). The neighbouring genes encoding hypothetical proteins were conserved between *w*Bol1J and *w*Bol1Y. The prophage insertion showed high homology to the MK-associated prophage region in *Wolbachia w*Hm-t and encoded the *Hb-oscar* gene (identical to *Hm-oscar*) as well as four *wmk* genes (identical to *wmk*-1 to *wmk*-4).

To clarify the genetic difference between the *w*Bol1 *Wolbachia* in the MK host line (Yogyakarta YOG-1 line, designated as *w*Bol1Y) and in the normal sex ratio host line (Ishigaki ISG-1 line, designated as *w*Bol1J), we completed closed genomes for both strains. The genomes of the two *w*Bol1 variants were generally colinear with a very high degree of similarity (ANI: 0.998) (**Fig. 2B**). However, the *w*Bol1Y genome carried a unique 63 kb insertion with a high degree of similarity to the MK-associated prophage WO*w*Hm-t76, found in the MK *Wolbachia w*Hm-t in the tortrix moth *Homona magnanima* (24) (**Fig. 2C**). Based on phylogenetic and molecular clock analyses of *Wolbachia* (41, 42), *w*Bol1J and *w*Bol1Y shared a common ancestor around 260-390 years ago. *w*Hm-t and the *w*Bol1 variants were estimated to have diverged approximately 118000-178000 years ago, indicating that the prophage region was acquired through lateral transfer.

We next examined the predicted coding content of the prophage acquired by *w*Bol1Y. The prophage region found in *H. magnanima*, WO*w*Hm-t76, encodes four *wmk* genes (*wmk-*1 to *wmk*-4) and the *Hm-oscar* gene, which confer highly penetrant male lethality when expressed in *Drosophila melanogaster* and *H. magnanima*, respectively (16, 24). Remarkably, genes identical at the nucleotide sequence level to the four *wmk* and the *Hm-oscar* (hereafter *Hb-oscar*) were conserved in the *w*Bol1Y-specific insertion (**Fig. 2C**, **Table S2**). *w*Bol1J and *w*Bol1Y carry an additional pair of *wmk*-3 and *wmk*-4 in another prophage region (**Fig. S1**), but *wmk*-1, *wmk*-2, and *Hb-oscar* were specific to *w*Bol1Y. *w*Bol1J and *w*Bol1Y also shared two pairs of *cif* genes (*cifA* and *cifB*) that were identical at the nucleotide sequence level, while *w*Bol1Y additionally carried a third *cifA* and *cifB* gene pair (similarly identical to the shared *cif* genes between *w*Bol1J and *w*Bol1Y) (**Table S2**).

We further compared the genome of *w*Bol1J and *w*Bol1Y to a *w*Bol1 variant from an MK host line collected from Rurutu, French Polynesia in 2004 (ERR12387130), and to the draft genome of an MK strain *w*Bol1b from Moorea, French Polynesia (43). The Rurutu MK strain was most similar to *w*Bol1J and lacked both the prophage insertion and *Hb-oscar* (*w*Bol1 Rurutu vs *w*Bol1J: ANI =0.998; vs *w*Bol1Y: ANI = 0.997) (**Fig. S2**). These data are consistent with *w*Bol1J as a suppressed MK with an *oscar*-independent MK mechanism. The draft genome from the Moorea MK strain was most similar to *w*Bol1Y (*w*Bol1 Moorea vs *w*Bol1Y: ANI =0.999; vs *w*Bol1J: ANI = 0.997). While the fragmented Moorea *w*Bol1b genome does not allow definitive proof of the presence or absence of *Hb-oscar*, BLAST searches identified two contigs with perfect matches to the *Hb-oscar* gene: CAOH01000067 (1435 bp) and CAOH01000068 (1535 bp), corresponding to *Hb-oscar* positions 327-1761 bp and 2012-3546 bp, respectively.

### Female-specific infection of *w*Bol1Y variant in *H. bolina*

PCR assays targeting the *Wolbachia wsp* gene revealed that all *Hypolimnas bolina* butterflies, both male and female, collected in Japan and Yogyakarta were infected with *w*Bol1 *Wolbachia* (**Fig. 3A**). Further PCR assays targeting *Hb-oscar* as a specific marker for *w*Bol1Y-type *Wolbachia* revealed that all Japanese female and male butterflies were negative for this marker gene. In contrast, in Yogyakarta, *Hb-oscar* was detected exclusively in females and in all the female-only producing (MK) butterfly lines (n = 13). In contrast, *w*Bol1-infected males from Yogyakarta were negative for the *Hb-oscar* gene (**Fig. 3A**).

**Fig. 3.**
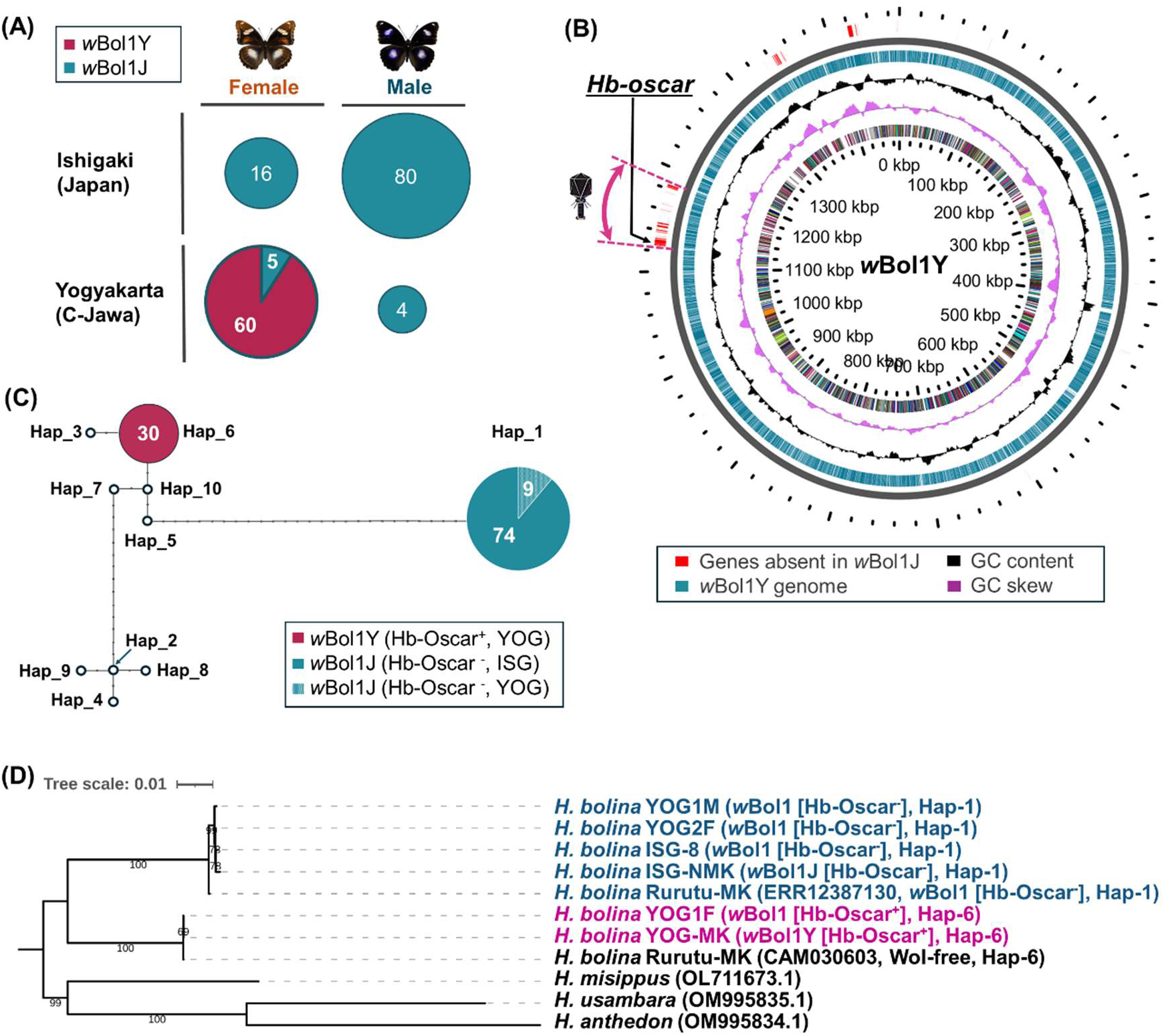
**Genome structures and prevalences of *w*Bol1 *Wolbachia* variants and their association with mitochondrial haplotypes.** (A) *Wolbachia* infection status of butterflies. *Wolbachia* infection was determined by amplification of the *Wolbachia*-specific *wsp* gene and Sanger sequencing. The genotype of *w*Bol1 (i.e. *w*Bol1Y [*Hb-oscar*-positive] or *w*Bol1J [*Hb-oscar*-negative]) was determined by PCR targeting the *Hb-oscar* gene. The numbers to the right of the pie charts indicate the number of individuals. (B) Comparative genomics of the *w*Bol1 genomes. Regions specific to *w*Bol1Y (absent in *w*Bol1J-type strains) are highlighted against the reference *w*Bol1Y genome. (C) Haplotype networks of the mitochondrial COI gene of *H. bolina* using *w*Bol1Y-infected females (n = 30 in Yogyakarta), *w*Bol1J-infected females (n = 12 in Ishigaki; n = 4 in Yogyakarta) and males (n = 62 in Ishigaki; n = 5 in Yogyakarta) collected in this study. Haplotype numbers were based on Charlat et al. (2009); we did not identify individuals carrying haplotype 2, 3, 4, 5, 7, 8, 9 and 10 in this study. (D) A phylogenetic tree based on the complete mitochondrial genomes of *Hypolimnas* species. *H. bolina* infected with *Hb-oscar-*bearing *w*Bol1Y (magenta) and *Hb-oscar*-deficient *w*Bol1J (blue).

We re-sequenced *w*Bol1-infected male and female butterflies collected in Japan and Yogyakarta using DNBSEQ short reads (Japan: 1 female and 1 male, both *Hb-oscar* negative; Yogyakarta: 2 females [1: *Hb-oscar* positive; 1: *Hb-oscar* negative] and 1 male, *Hb-oscar* negative). The genome of *Hb-oscar*-positive *w*Bol1 from a female, sampled in Yogyakarta, matched the reference *w*Bol1Y genome and carried the *w*Bol1Y-specific insertion. By contrast, *Hb-oscar*-deficient *w*Bol1 in a male and an unmated female sampled in Yogyakarta resembled the *w*Bol1J-type genome, lacking genes associated with the *w*Bol1Y-specific insertion including *Hb-oscar* (**Fig. 3B**). Similarly, the *Hb-oscar*-deficient *w*Bol1-infected Japanese female and male exhibited the *w*Bol1J-type genome. These results indicate the coexistence of *w*Bol1Y and *w*Bol1J-type *Wolbachia* within the Yogyakarta population, with distinct infection patterns between the sexes (*w*Bol1Y in females only, exhibiting MK; *w*Bol1J-type in both sexes, no MK). The presence of *w*Bol1J-type in both sexes is typical of a population where the suppressor of MK is at high frequency.

### Strong concordance between *w*Bol1Y and mitochondrial haplotype 6 in *H. bolina*

We analyzed the mitochondrial haplotypes of hosts infected with either *w*Bol1Y (bearing *Hb-oscar*) or *w*Bol1J (lacking *Hb-oscar*). Previous work established polymorphic sites within 414 bases of the mitochondrial COI gene that distinguished 10 different *H. bolina* mitochondrial haplotypes (37). *w*Bol1 has been identified from *H. bolina* carrying haplotypes 1, 3 and 6, and the suppressed MK *w*Bol1 is tightly associated with haplotype 1 (37, 38). In this study, hosts infected with *Hb-oscar*-deficient *w*Bol1, irrespective of their geographical origin, exhibited haplotype 1 (AJ844908.2) (Ishigaki: n = 73; Yogyakarta: n = 9). In contrast, all *w*Bol1Y-infected females were associated with haplotype 6 (n = 30, all from Yogyakarta) (**Fig. 3C**). We further analysed the phylogeny of *H. bolina* using their complete mitochondrial sequences (ca. 15 kb). The *w*Bol1Y-infected butterflies carrying haplotype 6 and *w*Bol1J-infected hosts carrying haplotype 1 were divided into two distinct clades (**Fig. 3D**). A *Wolbachia*-uninfected female carrying the haplotype 6, collected in Rurutu, French Polynesia (CAM035603) (44), was also grouped within the same clade as the *w*Bol1Y-infected *H. bolina* butterflies. In contrast, the female from the *w*Bol1J-infected MK Rurutu line (ERR12387130) exhibited haplotype 1.

### *w*Bol1Y is insensitive to the MK suppressor of *H. bolina*

The presence of *w*Bol1J-type (*Hb-oscar* deficient *w*Bol1) infection in male butterflies from Yogyakarta indicates the presence of the MK suppressor in this region, as males very rarely survive infection with MK *Wolbachia* in the absence of the suppressor (12). The persistence of the MK phenotype in the *w*Bol1Y-dominated host population is consistent with insensitivity to the MK suppressor. To determine the suppressor’s function against *w*Bol1Y-induced MK, we assessed the sex ratio of offspring from crosses between *w*Bol1J-infected males (carrying the suppressor) and *w*Bol1Y-infected females. Through introgression, nuclear genetic material from the *w*Bol1J-bearing male (with the suppressor) is matched to the *w*Bol1Y infection in the hybrid offspring (as *Wolbachia* is transmitted maternally).

Previous work demonstrated that the suppressor mutation against *w*Bol1-induced MK is dominant, as male offspring survival was restored in one generation of crosses between females from MK lines (suppressor-negative/*w*Bol1 infected) and suppressor-bearing *w*Bol1-infected males (non-MK) (12). In contrast to previous crosses, the progeny of *w*Bol1Y-infected females and *w*Bol1J-infected males were all female (122 females vs. 0 males; P = 0.000, binomial test), demonstrating a lack of suppression, and contrasting with a cross between *w*Bol1J-bearing male and female (27 females vs. 30 males; P = 0.396) (**Fig. 4**). The egg hatchability of *w*Bol1Y-infected lines was approximately 50%, indicating male death during embryogenesis. These data also indicates that there is no cytoplasmic incompatibility (CI) between *w*Bol1J infected males and *w*Bol1Y infected females, and that the strains share a compatibility type (45).

**Fig. 4.**
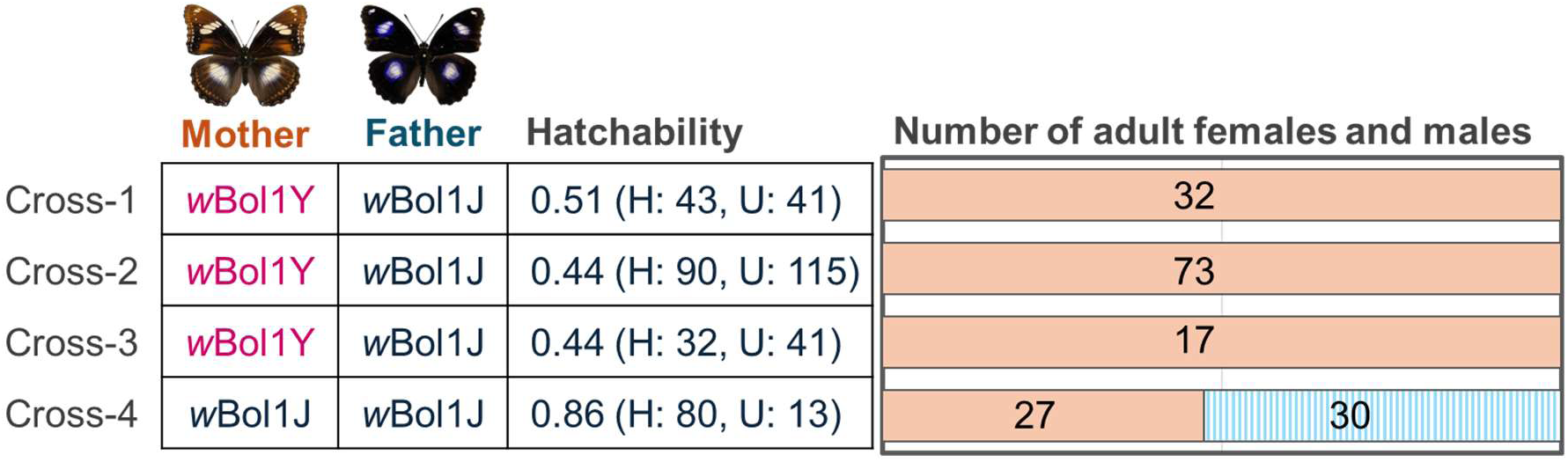
**Sex ratio in the crosses of *w*Bol1Y-bearing females and *w*Bol1J-bearing males with suppressor.** The genotype of mothers and fathers was determined by detection of *Wolbachia wsp* and the *Hb-oscar* gene (*Hb-oscar* positive: *w*Bol1Y-type; *Hb-oscar* negative: *w*Bol1J-type). The hatchability of the offspring was calculated based on the numbers of hatched individuals (shown as H) and unhatched embryos (shown as U). Numbers on the bars indicate numbers of female (orange) and male (blue) individuals in the offspring.

### *w*Bol1Y induces female-type sex determination and impairs dosage compensation in male *H. bolina* embryos

We verified the effects of *Wolbachia* infection on the butterfly’s sex determination gene expression and splicing patterns. The lepidopteran sex determination system is a cascade where genes are differentially spliced in male and female fated embryos (46). The primary male sex determinant is the gene *masculinizer* (*masc*), which regulates both dosage compensation and the downstream sex determination cascades. The male signal initiated by the *masc* gene leads to the male-type splicing of the downstream sex determinant *doublesex* (*dsx*), which in the absence of this signal shows female-type splicing (47). The splicing pattern of the *dsx* gene is additionally regulated by *zinc finger protein 2* (*znf2*) in *Bombyx mori* (48). MK *Wolbachia* strains generally ‘feminise’ the sex determination cascades in males, with MK *Wolbachia*-infected hosts exhibiting female-type splicing variants of *masc*, *znf2*, and *dsx* genes *in vivo* and *in vitro* (17, 18, 22, 49, 50).

We first determined the splicing isoforms of *dsx* in *H. bolina* by analyzing RNA extracted from the abdomens of adult females and males, as well as from embryos. Adult female and male *H. bolina* expressed four and three splicing isoforms of *dsx*, respectively (**Fig. 5A**). Among these isoforms, *HbdsxF-1* was dominant in females, while *HbdsxM-1* was dominant in males. Notably, other isoforms were specifically expressed in the line derived from Ishigaki, Japan (ISG), which is infected with the suppressed variant *w*Bol1J. In *Wolbachia-*uninfected and *w*Bol1J-infected/suppressor positive *H. bolina* embryos, females and males were identified by the presence and absence of W chromatin. Female embryos exhibited *HbdsxF-1* splice form and male embryos *HbdsxM-1*. Contrastingly, *w*Bol1Y-infected *H. bolina* embryos induced female-type *dsx* splicing (*HbdsxF-1*) regardless of the presence or absence of W chromatin (**Fig. 5B**). *w*Bol1Y-infected male embryos also exhibited weak male-type *dsx* splicing (*HbdsxM-1*) in addition to the female-type splicing, similar to the previous observation in *Homona* male embryos infected with *w*Hm-t or expressing *Hm-oscar* (16, 17).

**Fig. 5.**
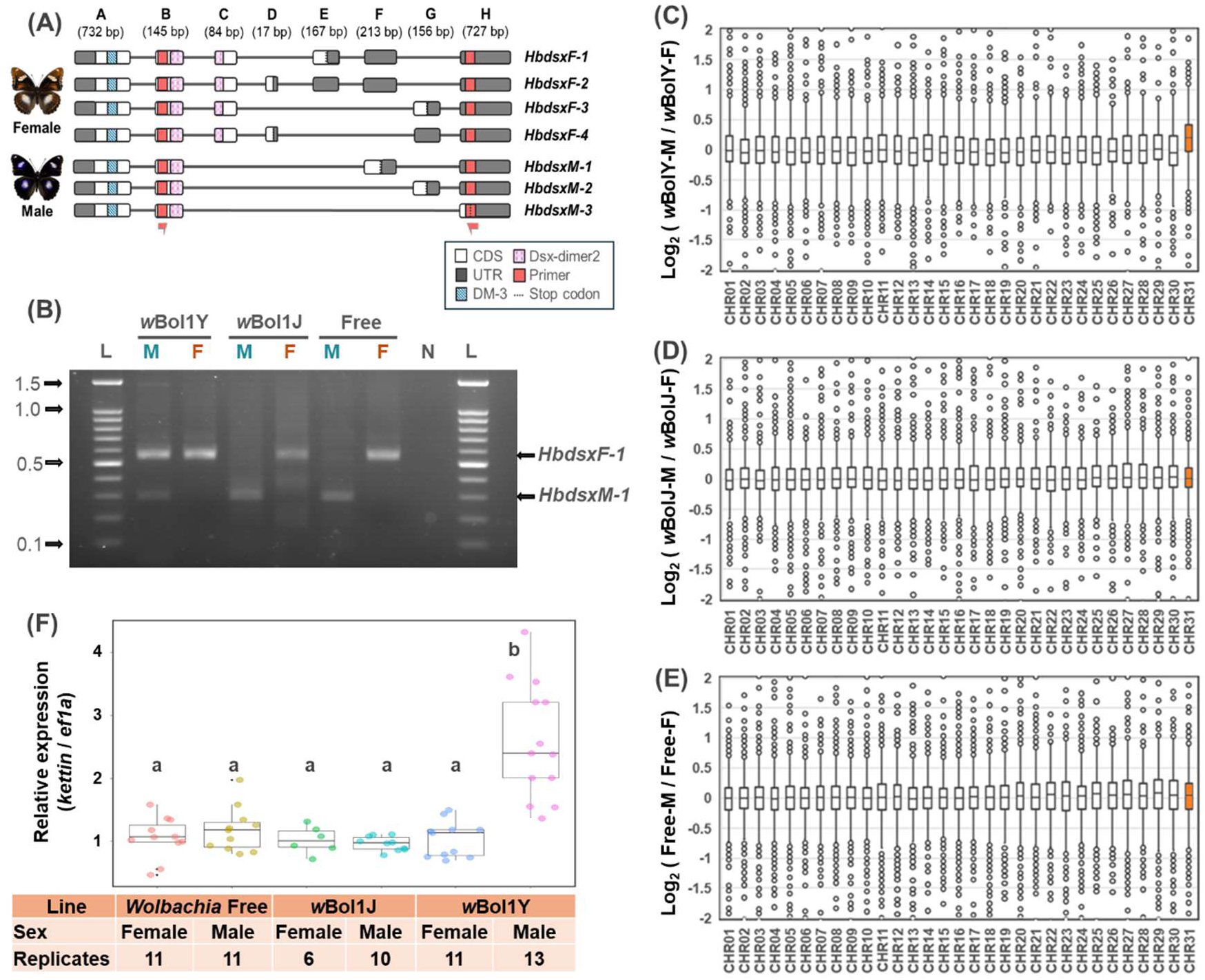
**Effects of *w*Bol1 *Wolbachia* on dosage compensation and sex determination systems in *H. bolina* embryos** (A) Splicing isoforms of *Hbdsx*. ISG line infected with *w*Bol1J expressed additional isoforms (*HbdsxF-2-4* in females and *HbdsxM-2-3* in males) that were not observed in *w*Bol1Y-infected or uninfected host lines. (B) Splicing patterns of the downstream sex-determining gene *dsx* in *H. bolina* embryos. M and F indicate W chromatin-negative (ZZ: male genotype) and W chromatin-positive (ZW: female genotype) mature embryos, respectively. *w*Bol1Y-infected, *w*Bol1J-infected, and uninfected males predominantly expressed the dominant isoform *HbdsxM-1*, as shown in panel E. *w*Bol1Y-infected males additionally expressed the dominant isoform of *HbdsxF-1*, which is typically observed in *w*Bol1Y-infected, *w*Bol1J-infected, and uninfected females. L: 100 bp DNA ladder (ExcelBand 100 bp DNA Ladder, SMOBIO Technology, Inc., Hsinchu, Taiwan). 0.1, 0.5, 1.0, and 1.5 kb markers are indicated with arrows. N: negative control (water). (C-E) Normalized expression levels (TPM) and chromosomal distributions of transcripts in *H. bolina* embryos. RNA-seq data of embryos (108 hpo) were used to make the following comparisons: *w*Bol1Y-infected males versus females (C), *w*Bol1J-infected males versus females (D), and *Wolbachia*-negative males versus females (E). The X axis represents the chromosome number of *H. bolina*. (F) Relative expression of a Z-linked *kettin* gene in *H. bolina* groups. Different letters indicate statistical significance (Steel-Dwass test, P < 0.05).

The impact of *Wolbachia* infection on dosage compensation in *H. bolina* was then assessed by quantifying the expression of Z chromosome-linked genes. In Lepidoptera, expression levels of the Z-linked genes are balanced by dosage compensation between males (with ZZ karyotype) and females (with ZW). RNA-seq analysis using the *H. bolina* genome (44) as the reference, revealed that Z-linked genes in *w*Bol1Y-infected embryos had higher expression in males than in females (**Fig. 5C**), which was not observed in the *Wolbachia-*uninfected and the suppressed *w*Bol1J-infected groups (Z=chromosome 31 [CHR31], **Fig. 5D-E**). The high (i.e. doubled) Z-linked gene expression in *w*Bol1Y-infected males was further confirmed by quantification of the expression of the Z-linked *kettin* gene (**Fig. 5F**; all statistical data are provided in **Table S3**). These data indicate that unsuppressed *w*Bol1Y impairs the host’s dosage compensation system, mirroring the MK mechanism observed in other Lepidoptera (17, 18)

### Cell-based assays demonstrate that suppressed *w*Bol1J and unsuppressed *w*Bol1Y alter expression of lepidopteran sex determination cascades

The impacts of the suppressed and unsuppressed *w*Bol1 variants on lepidopteran sex determination systems were further investigated using a lepidopteran cell culture system. For this assay, we employed the recently established cell culture system OsM1, derived from male moths of *Ostrinia scapulalis* (ZZ karyotype) (50). The male-derived OsM1 cells transinfected with the *w*Bol1Y and *w*Bol1J progressively expressed female-type splicing variants for *dsx* (*OsdsxF*), *masc* (*OsmascF*), and *znf2* (*Osznf2F*) (**Fig. 6A-D**), in contrast to the male splicing pattern observed in control (uninfected) cells. Notably, the feminisation of the sex determination cascades by *Hb-oscar*-deficient *w*Bol1J was established later than for *Hb-oscar*-bearing *w*Bol1Y (*w*Bol1J: 12-16 weeks post transinfection [wpt]; *w*Bol1Y: 4-8 wpt). Thus, both the suppressed *w*Bol1J as well as the unsuppressed *w*Bol1Y retain the ability to inhibit the lepidopteran masculinizing pathway, as observed in other MK *Wolbachia* (17, 18, 22, 49).

**Fig. 6.**
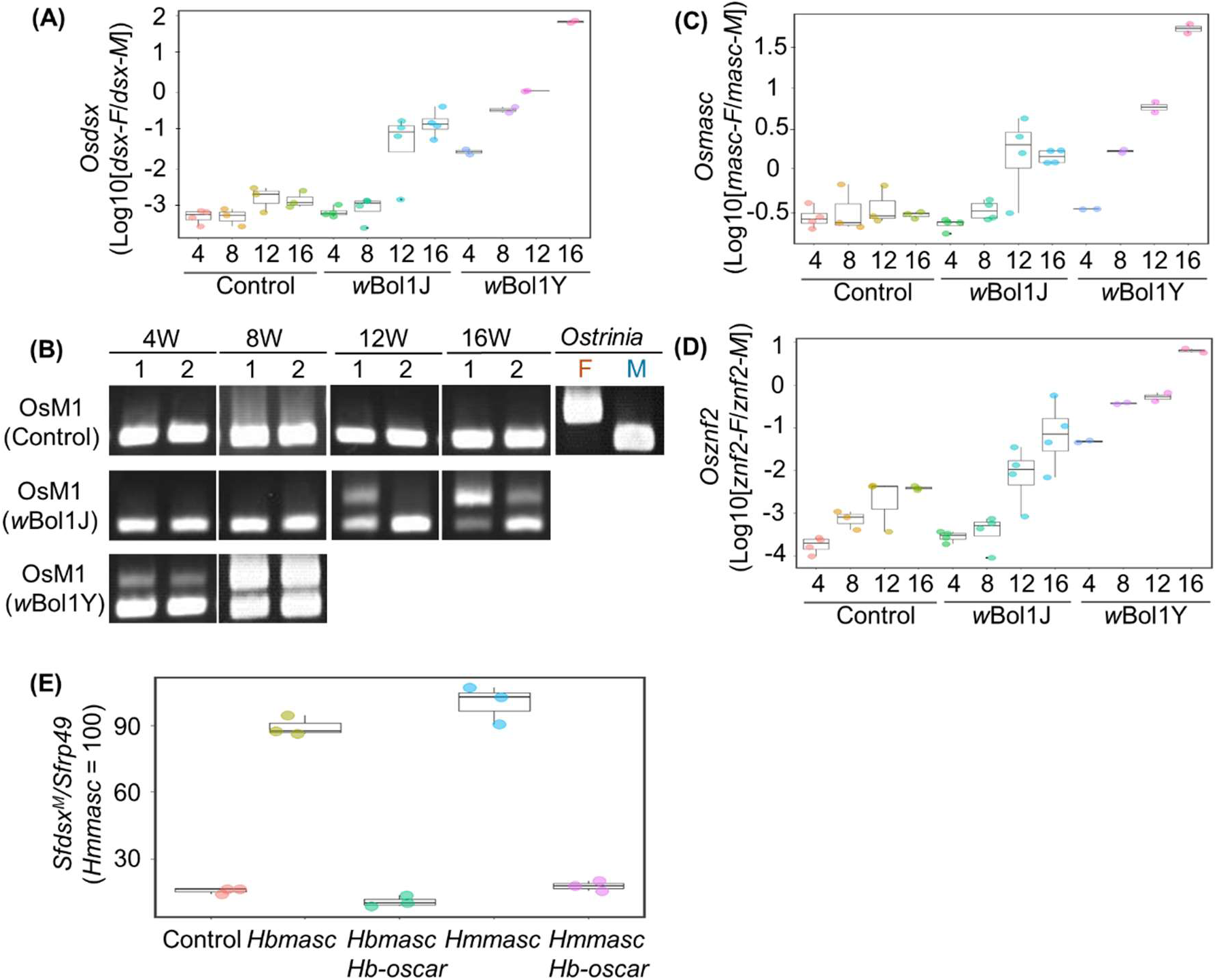
**Effects of *w*Bol1 *Wolbachia* and *Hb-oscar* on expression of lepidopteran sex determination systems *in vitro*** (A) The relative expression of *OsdsxF/OsdsxM* was quantified in *Wolbachia* transinfected and control cells. Cells were harvested every four weeks and x-axis indicates the time after *Wolbachia* transinfection (e.g. 12 weeks after transinfection). Dot plots indicate replicates. The splicing patterns of *dsx* were also verified by the diagnostic PCR assays. F: female; M: male (B). *OsmascF/OsmascM* (C) and *Osznf2F/Osznf2M* (D) were also quantified in *Wolbachia*-transinfected and control cells. Dot plots indicate replicates. (E) Relative expression levels of the male-specific *Sfdsx^M^* variant in transfected Sf9 cells. The relative expression of *Sfdsx^M^*in the control (non-inserted pIZ/V5) and *Hb-oscar* co-transfected groups was normalized by setting the mean in the *Hmmasc* single transfected group to 100 (n = 3, technical replicates in each condition). Similar results were obtained in two independent experiments (n = 3, technical replicates in each condition).

### *Hb-oscar* is sufficient for inducing feminization activity *in vitro*

The prophage element that is private to *w*Bol1Y carries a copy of the *Hb-oscar* gene, which represents a candidate additional MK element in the *w*Bol1Y genome. The *Hb-oscar* gene is identical to *Hm-oscar* gene that was previously shown to induce MK in the moth *H. magnanima* by disrupting the masculinizing function of the *masc* gene (*Hmmasc*) (16). To test whether *Hb-oscar* has such activity, we investigated whether it could suppress the function of the primary male determinant *masc* gene of *H. bolina* (*Hbmasc*).

We performed this analysis *in vitro* by transfection using Sf9 cells, which are derived from the female ovaries of *Spodoptera frugiperda* (51) and exhibit a female-type sex determination (20). As observed previously (20), transfection with the *Hbmasc*-inserted pIZ/V5-His vector induced male-type sex determination in Sf9 cells, as indicated by increased expression of the male-specific splicing isoform of the *dsx* gene (*dsx-M*) (**Fig. 6E**). In contrast, cells transfected with the non-inserted pIZ/V5-His vector exhibited standard female sex determination with a low expression level of *dsx-M*.

We then tested whether *Hb-oscar* could revert cells to female-type expression by suppressing the function of *masc* genes. Co-transfection of *Hbmasc/Hb-oscar*-inserted pIZ/V5-His vectors suppressed the masculinizing function of *Hbmasc* (**Fig. 6E**), evidenced by the lower expression levels of *dsx-M* compared to cells transfected with *Hbmasc* alone. Expression levels of *dsx-M* in the co-transfected cells were similar to control cells. Co-transfection of *Hb-oscar* also suppressed the masculinizing function of the *H. magnanima*-derived *Hmmasc*. These results support the hypothesis that *Hb-oscar* acts as an MK agent in *H. bolina*.

## Discussion

Sex ratio distorting *Wolbachia* alter the evolutionary ecology of their host in two ways – through altering the reproductive ecology of their host (mediated through the population sex ratio bias they produce) and through selection to prevent their action – suppression of MK. Previous work on the *H. bolina-w*Bol1 MK system in Samoa demonstrated that the male-killer *w*Bol1 could reach high prevalence and reduce fertility through paucity of males (29). However, the MK phenotype (and extreme sex ratio bias) disappeared rapidly with the spread of a single locus suppressor (10, 12, 52, 53).

In the present study, we present a similar extreme sex ratio bias in the species in Yogyakarta, Jawa Island, Indonesia, also associated with high prevalence of a *w*Bol1 MK *Wolbachia*, and loss of female fertility in the population through paucity of males. The population contrasts with those elsewhere in Southeast Asia, including the adjacent island of Borneo (Saba), where extreme sex ratios associated with *w*Bol1-induced MK have been recorded historically, but have disappeared following spread of the suppressor of MK (37, 38). Indeed, the suppressor presence in Yogyakarta *H. bolina* is indicated by the presence of male butterflies that carry the non-*Hb-oscar* carrying version of *w*Bol1 (*w*Bol1J-type). The MK trait in the Yogyakarta population was insensitive to the MK suppressor that evolved against the MK *w*Bol1. Genomic analyses showed the MK strain in Yogyakarta, *w*Bol1Y, was very closely related to, but showed key genetic differences from, the ‘suppressed’ *w*Bol1. Notably, *w*Bol1Y carried an extra prophage element that contained strong candidate genes for MK.

The presence of an MK variant that is insensitive to the circulating MK suppressor is significant for two reasons. First, our observations indicate that multiple MK mechanisms can evolve within a single host species. While MK mechanisms are known to be diverse amongst symbionts and between host species (16, 19, 20, 24, 25, 54–56), we observed two closely related variants, with distinct MK phenotypes (suppressible/not suppressible) circulating within a population. Second, the presence of a MK variant that is insensitive to the circulating MK suppressor is also consistent with the model of an escalating arms race. Rather than male-killer presence giving rise to selection for suppression and then an end point of loss of the MK phenotype, our data indicate insensitivity to suppression is an evolvable trait. Thus, suppression of MK is not necessarily an endpoint, and the dynamics of suppression and insensitivity to suppression are likely to be cyclical.

### Evolution of MK mechanism within a host species

The diversity of MK mechanisms has been considered to relate to diversity in insect sex determination systems, with MK a convergently evolved symbiont phenotype underpinned by diverse genetic mechanisms ‘fitting’ the specific sex determination and dosage compensation systems of host species (16, 20, 24, 55, 56). Our study reflects diversity of MK mechanism/system within a host species. Whilst previous work had established multiple MK strains from different bacterial groups/strains circulating within a host species (17, 57, 58), the very close relatedness of strains with distinct MK phenotypes in our study is most parsimoniously explained by one evolving from the other within the host species. The evidence for this is fivefold. First, *w*Bol1Y has distinct properties in relation to MK, notably it is insensitive to the MK suppressor that silences other *w*Bol1 variants. Second, *w*Bol1Y clearly shares a recent common ancestor with the suppressor-sensitive MK strain, with very high ANI identity indicating a common ancestor estimated in the last 230-390 years. Third, *w*Bol1Y is distinct from the suppressed strain in carrying a prophage element harboring genes involved in MK in other lepidopteran systems (e.g. *Hb-oscar*). This gain of function is most parsimoniously explained through the evolution of *w*Bol1Y from *w*Bol1. Fourth, *w*Bol1Y induces female-specific splicing of *dsx* and impairment of dosage compensation in male *H. bolina* embryos and we observed feminization of male *Ostrinia* cells *in vitro*. This feminization was observed to occur more rapidly for *w*Bol1Y than the suppressible *w*Bol1J isolate tested. Fifth, *Hb-oscar,* unique to the unsuppressed *w*Bol1Y, impairs the masculinizing function of the *H. bolina masc* gene *in vitro*. These data indicate *w*Bol1Y is a form of *w*Bol1 with additional MK factors, and evolution within a host species to alter the MK mechanism.

Differences in suppressor sensitivity likely arise from distinct MK mechanisms in *w*Bol1Y and *w*Bol1J. *Hb-oscar*, located on the unique prophage insert in *w*Bol1Y, is itself sufficient for feminization *in vitro* and is a strong candidate for being a factor insensitive to suppression. The prophage also carries additional candidate MK factors whose functions has yet to be determined (*wmk*-1/*wmk*-2 pair and an additional *wmk*-3/*wmk*-4 pair). *w*Bol1Y likely achieves MK by disrupting the dosage compensation system by ablating the function of *masc* gene, which is responsible for regulating both dosage compensation and downstream sex determination. This mechanism is consistent with observations in *Ostrinia* and *Homona* moths infected with the *oscar*-bearing *Wolbachia* (16, 20).

Given that the MK suppressor does not silence *w*Bol1Y-induced MK, the suppressor likely interacts with as yet-uncharacterized MK factor(s) in *w*Bol1J. Whilst the suppressed *w*Bol1J-induced MK mechanism in the host remains to be determined, we demonstrated through the cell culture system that *w*Bol1J feminizes the sex determination system of *Ostrinia* males. This finding is consistent with the observation that certain *oscar*-deficient *Wolbachia* induce female sex determination in the cell system (49). While the effects in native hosts may differ from those in *Ostrinia* cells, *w*Bol1J appears to target the host sex determination system to induce MK in *Hypolimnas* via a mechanism distinct from *w*Bol1Y.

### Evolution of male killers, host suppression and escalating arms races

The evolutionary arms race between selfish sex-ratio distorters and counteracting suppressor genes can drive rapid flux in sex ratios, manifesting as escalating cycles of selection between host and parasitic elements (27, 59, 60). Escalation has been characterized in the interaction between meiotic drive systems and suppressors of drive (61), and for mitochondria inducing cytoplasmic male sterility and nuclear restorer genes (62–64). In these cases, the escalation is considered an important contributor to the generation of hybrid sterility between populations (61). MK symbionts likewise drive host evolution as antagonists to be avoided because MK imposes a fitness cost on their host (26). There have been numerous records of MK circulating in populations, and more recently broad evidence of suppression of MK (10–15). However, in contrast to drive systems, counter-selection on the symbiont to escape suppression has not yet been evidenced. A key requirement for an escalating arms race—and the potential for coevolutionary cycles—is that suppressed MK strains can evolve to become insensitive to suppression.

Our data present a potential evolutionary pathway in *H. bolina*, in which the spread of MK is followed by the suppressor spread, subsequently driving the evolution of a novel non-suppressible form of MK, *w*Bol1Y. In this model, the insensitive MK form is an evolved response to suppression. Our hypothesis is that non-suppressibility derives from the inserted prophage material that includes additional MK factors, and the action of these factors is unaffected by the suppressor. The suppressed *w*Bol1 is strongly linked to mitochondrial haplotype 1, while unsuppressed *w*Bol1Y was associated with haplotype 6. The *w*Bol1 variants likely evolved within distinct host lineages, with *w*Bol1Y acquiring an MK-associated prophage region similar to that found in a distantly related MK strain *w*Hm-t. Phage movement/integration may have played a crucial role in re-establishing MK in *w*Bol1Y, paralleling the evolution of the MK phenotype in *w*Hm-t from the closely related non-MK *w*Hm-c in *Homona* moths (24, 65).

*w*Bol1J and the additional prophage-bearing *w*Bol1Y, are very closely related with an estimated time of divergence, from ANI, of 260-390 years ago. Previous analysis of museum specimens showed that suppression of *w*Bol1-induced MK in the Philippines was present in 1894 (130 years ago), the earliest specimens tested from the island (38). However, there is insufficient evidence to establish whether the unsuppressed *w*Bol1Y form arose as a counter-reaction to the spread of the suppressor, or whether it was pre-existing and then retained by virtue of its capacity to avoid suppression. Further investigation on *Wolbachia* genomes and suppressor locus using historical and current *H. bolina* samples collected broadly would be necessary to resolve this question.

## Materials and methods

### Insects

*H. bolina* adults were collected in Ishigaki (Japan) and Yogyakarta (Indonesia). The sex of the butterflies was determined based on the color and spot patterns on their wings. The captured adults were reared on ion-supplemented water Pocari Sweat (Otsuka Pharmaceutical, Tokyo, Japan) and subjected to egg collection without mating after capture. For oviposition, butterflies were placed in a plastic cup (90 mm diameter, 120 mm in hight) with food plants (*Ipomoea batatas*, *Ipomoea aquatica*, and *Achyranthes bidentata*). The eggs oviposited on the plants were collected every day until the female died. The hatched larvae were reared on the food plants until pupation. The emerged adults were sexed based on their color patterns and subjected to egg collection or the downstream assays. To obtain eggs from the laboratory reared lines, 10 males and 10 females were placed in a cage (5m x 5m x 5m) for mating in the daytime (AM 7-PM 4), and the mated females were used for egg collection. Alternatively, hand-paired females were used. All insects were reared at a 16L:8D light cycle, at 25°C, and with 60% relative humidity.

### Hatchability and sexing of the embryos

The hatchability of each butterfly line was calculated by counting the number of hatched and unhatched embryos four days after the first individual hatched (typically 8 days after oviposition). The hatched neonates (1^st^ instar larvae) and unhatched embryos were sexed by observing the female-specific heterochromatin (W chromosome) under microscopy as described previously (33, 66) with some modifications. Briefly, neonates or unhatched late-stage embryos were dissected on glass slides with forceps. Malpighian tubules were fixed with methanol/acetic acid (50% v/v) and stained with lactic acetic orcein for W chromatin observations (presence: female; absence: male).

### Antibiotic treatment

Antibiotic treatment was performed as described in Mitsuhashi et al. (32) with several modifications. Briefly, mated female adults were fed with the 0.2% (W/W) rifampicin-supplied Pocari Sweat (Otsuka Pharmaceutical) for six days. As a control, females from the same isofemale line were fed with an antibiotics-free Pocari Sweat (Otsuka Pharmaceutical). Eggs were collected every day until the females died. The larvae were reared as mentioned above, and individuals collected on days 1 to 6 of rifampicin treatment were reared separately. The sex ratio of adults in each daily group was recorded separately and sex was determined by wing color pattern.

### *Wolbachia* and other endosymbionts detections

Total DNA was extracted from the leg or abdomen of the butterflies using DNeasy Blood & Tissue Kits (Qiagen, Hyden, Germany) following the manufacture’s protocol. About 50-100ng total DNA was used for PCR assays targeting *Wolbachia wsp* and MLST genes (*ftsZ, gatB, coxA, hcpA,* and *fbpA*) (**Table S4**) using EmeraldAmp PCR Master Mix (TaKaRa Bio Inc., Shiga, Japan). The mitochondrial COI gene was also amplified and used as a control (**Table S4**). The amplification reaction parameters were as follows: 3 min at 94 °C; followed by 35 cycles of 30 sec at 94 °C, 30 sec at 60 °C, and 30 sec at 72 °C; with a final extension for 7 min at 72 °C. The amplicons were sequenced using sanger sequencer as described previously (67).

To check for infection with other potential male killers in the butterfly lines, we performed 16s ribosomal metagenomic sequencing. Briefly, the DNA extracted were amplified with primers targeting bacterial the hypervariable 16S rRNA V3-V4 region (**Table S4**). The libraries generated from the amplicons were sequenced using MiSeq with MiSeq Reagent Kit v3 (Illumina, CA, USA) for 300 bp paired end reads. The obtained sequence data were trimmed with FASTX-Toolkit v0.0.14 (https://github.com/agordon/fastx_toolkit) and sickle v1.33 (https://github.com/najoshi/sickle) to remove low quality reads (Q score below 20; sequence length below130 bp). The paired end reads were merged using FLASH v1.2.11 (68) and were analysed using Qiime2 v2022.8 (69). The taxa of the bacteria were annotated using the Silva v.138 database (70).

### Genome analyses

High molecular weight DNA was extracted from the ovaries of *Wolbachia*-infected female adults following the manufacture’s protocol of the NanoBind Big DNA Tissue kit (Circulomics, Baltimore, MD, USA). The extracted DNA was sequenced using DNBSEQ-G400 with paired-end 150 bp (MGI tech, Shenzhen, China) and MinION with ligation sequencing kit for MinION flow cell [R10.4] (Oxford Nanopore, Oxford, UK). The Nanopore data were assembled using Flye v1.6 (71). Subsequently, *Wolbachia* contigs were extracted from the assembled contigs after a series of Blast searches. The extracted *Wolbachia* genomes were polished 3–5 times with Illumina data using Pilon (72) and minimap2 (73). The resulting polished *Wolbachia* genomes were annotated using DFAST (74).

A dot-plot of the genome structures of *w*Bol1 *Wolbachia* variants were visualised using D-Genies (https://dgenies.toulouse.inra.fr/). Average nucleotide identity (ANI) of *Wolbachia* genomes were calculated using pyani (https://github.com/widdowquinn/pyani). Prophage regions were annotated using the PHASTEST web server (https://phastest.ca/). In addition, the MK-associated prophage WO*w*Hm-t76 region of the MK *Wolbachia w*Hm-t in *H. magnanima* was searched in the *w*Bol1 genomes by Blastn. Homologies of the WO*w*Hm-t76 region and other WO phages were visualized using Easyfig v2 (75) and TBtools (76). To check the presence of the phenotype-associated *Wolbachia* genes, *w*Bol1Y and *w*Bol1J-encoded genes were blasted against the homologs of *oscar* (20, 24, 49), *wmk* (24, 25), and *cif* (77).

Field-collected butterflies from Yogyakarta (2 females and a male) and Japan (a female and male) were sequenced using DNBSEQ-G400 with paired-end 150 bp (MGI tech). The sequence data were then mapped onto the *w*Bol1Y genome using minimap2 (73), and the consensus genomes were extracted using SAMtools (78). Syntenies of the *w*Bol1 genomes were visualized using GView web server (https://server.gview.ca/).

### Phylogenetic analysis

MLST (*gatB, coxA, hcpA, ftsZ*, and *fbpA* genes) and *wsp* fragments from the *w*Bol1Y and *w*Bol1J strains were aligned using ClustalW (79) with a those of *Wolbachia* endosymbionts reported previously (57, 80). In addition, single-copy genes conserved among *Wolbachia* strains were obtained using OrthoFinder (81) and concatenated, aligned, and trimmed using SeqKit (82), MAFFT (83), and trimAl (84), respectively. These alignment files were used to construct a phylogenetic tree based on maximum likelihood with bootstrap resampling of 1000 replicates using the IQTREE server (http://iqtree.cibiv.univie.ac.at/) and visualized using the iTOL web server (https://itol.embl.de/). Haplotype network analysis was conducted on mitochondrial COI sequences of *H. bolina* using DnaSP6 (85) and Network (86), as described in Arai et al (65).

Molecular clock analysis of *w*Bol1 variants and the *w*Hm-t strain was performed using branch lengths of nodes on the phylogenetic tree derived from their single-copy orthologs. Following the estimates by Richardson et al. (41) and Schorz et al. (42), we applied an annual substitution rate of 0.8–1.2 substitutions per million years.

### Detection of the *Hb-oscar* gene

DNA extracted from the *H. bolina* samples was subjected to PCR using primer sets, as described by Arai et al. (24) **(Table S4)**. Briefly, *Hb-oscar* was amplified using Emerald Amp Max Master Mix (TaKaRa) at 94°C for 3 min; 35 cycles of 94°C for 30 s, 62°C for 30 s, and 72°C for 3 min; and a final extension at 72°C for 7 min.

For qPCR, 10 ng DNA was amplified with a primer set and a TaqMan probe (**Table S4**) with PrimeTime Gene Expression Master Mix (Integrated DNA Technologies IDT, NJ, USA). For 10 μl reaction, 5 μl PrimeTime Gene Expression Master Mix, 0.5 μl forward primer (10 μM), 0.5 μl reverse primer (10 μM), 0.2 μl probe (10 μM), and 2.8 μl UltraPure™ DNase/RNase-Free Distilled Water (Invitrogen, MA, USA) were mixed with 1 μl DNA (10 ng/μl). The PCR reaction was performed using LightCycler® 96 (Roche, Basel, Switzerland) with the following reaction: 95°C for 3 min; 45 cycles of 95°C for 15 s and 60°C for 1 m.

### Crossing experiment

Unmated *w*Bol1Y-infected or *w*Bol1J-infected females were mated with *w*Bol1J-infected males in a cage or by hand pairing as mentioned above. The resulting mated females were subjected to egg collection, and the larvae hatched from eggs collected were reared on the host plants. The sex of eclosed adults were determined based on their wing color patterns, and the sex ratio of broods was calculated.

### RNA extraction from adults and embryos

RNA was extracted from the *w*Bol1Y-infected, *w*Bol1J-infected, and *w*Bol1Y-cured (i.e., uninfected) *H. bolina* lines. For adults, male and female butterflies were frozen at −80°C, and whole abdomens were subjected for RNA extraction. Mature embryos showing black head capsule (96 hour post oviposition, 25°C) were dissected on glass slides using forceps, as described in Arai et al. (17). Their Malpighian tubules were fixed with methanol/acetic acid (50% v/v) and stained with lactic acetic orcein for W chromatin observations. The remaining tissues not used for sexing were stored in ISOGEN II (Nippon Gene) at −80°C until subsequent extraction. Adult abdomens and sex determined embryos (confirmed based on the presence or absence of W chromatin) were homogenized in ISOGEN II to extract RNA as described in Arai et al.(17). In brief, 0.4 times the volume of UltraPure distilled water (Invitrogen) was added to the ISOGEN II homogenates, which were centrifuged at 12000 × *g* for 15 min at 4°C to pellet proteins and DNAs. The resulting supernatant was mixed with the same volume of isopropanol to precipitate RNAs; then, the resulting solutions were centrifuged at 20,400 × *g* and 4°C for 60 min. The RNA pellets were washed with 80% ethanol and were centrifuged at 20,400 × *g* and 4°C for 15 min. After removing all supernatant, the pellets were dried for 15 min at room temperature and eluted in 10 µL of UltraPure distilled water (Invitrogen).

### RNA-seq and homology surveys

A total of 1.0 µg of RNA (an individual for adults; 3-5 pooled individuals for embryos) from *w*Bol1J, *w*Bol1Y, and uninfected butterflies and embryos were used to prepare mRNA-seq libraries via the NEBNext Poly (A) mRNA Magnetic Isolation Module (New England Biolabs) and the NEBNext Ultra II RNA Library Prep kit for Illumina (New England Biolabs) following the manufacturer’s protocol. The adaptor sequences and low-quality reads (Qscore <20) were removed from the generated sequence data [150 bp paired-end (PE150)] using Trimmomatic (87). The trimmed data from adults were pooled and *de novo* assembled using Trinity (88). Homologies between the assembled transcripts and known lepidopteran *masc*, *dsx*, *kettin* (Z-linked gene), and *ef1a* (autosome control gene) genes were analyzed using Blastn. Coding sequences of these genes were predicted using ORFfinder (https://www.ncbi.nlm.nih.gov/orffinder/).

### *Hbdsx* detection

Based on the sequences of contigs that showed high homology to the known *dsx* homologs in Lepidoptera, we designed a primer pair amplifying first and last exons for *Hbdsx* shared by both males and females. We determined the splicing isoforms of the *Hbdsx* as described in Arai et al. (17) with some modifications. In brief, total RNA (100– 300 ng) extracted from sex-determined mature embryos/adults was reverse-transcribed using PrimeScript™ II Reverse Transcriptase (TaKaRa, Shiga, Japan) as described above. Then, cDNA was amplified using KOD-FX Neo (Toyobo Co., Ltd.) with the primers Hbdsx.Ex1.503-522F1 and Hbdsx.Ex7.40-60R1 (**Table S4**). The PCR conditions used were as follows: 94°C for 2 min, followed by 45 cycles of 98°C for 10 s, 60°C for 30 s, and 68°C for 30 s. The amplicons were cloned into pGemT vector and sanger sequenced by using T7 and SP6 primers at Eurofins Genomics. Based on the sequence data of *dsx* isoforms in the *w*Bol1Y/*w*Bol1J-infected hosts, we designed a new primer to distinguish male and female *dsx* isoforms: HbExon3.123F and HbExon7.51-70R. PCR was performed at the same condition as mentioned above, and amplicons were subjected to electrophoresis on 1.5% agarose Tris-borate-EDTA buffer (89 mM Tris-borate, 89 mM boric acid, 2 mM EDTA) gels.

### Quantification of Z chromosome-linked genes

The trimmed RNA-seq reads derived from *w*Bol1Y-infected, *w*Bol1J-infected, and *w*Bol1Y-cured (uninfected) male and female embryos were mapped to the previously assembled genome data (transcripts data) for *H. bolina* (44) using Kallisto (89) to generate the normalized read count data (transcripts per million, TPM). The binary logarithms of the TPM differences between males and females belonging to each *H. magnanima* group were calculated to assess the fold-changes in gene expression levels between sexes. The binary logarithms of the TPM differences between males and females in the *H. bolina* chromosomes 1 to 31 were plotted (chromosome 31 represents Z chromosome).

To confirm the effects observed in the RNA-seq-based quantification of the Z chromosome gene, expression level of the Z-linked *Hbkettin* gene was quantified by reverse transcription-qPCR. Briefly, total RNA (100–300 ng) extracted from an embryo was reverse transcribed using PrimeScript II reverse transcriptase (TaKaRa) at 30°C for 10 min, 45°C for 60 min, and 70°C for 15 min. The cDNA was diluted 10 times by water and used to quantify relative gene expression levels, with normalization to the control gene elongation factor 1a (*ef1a*). qPCR was performed using primer sets for *Hbkettin* and *Hbef1a* (**Table S4**), and KOD SYBR qPCR Mix (Toyobo, Osaka, Japan) in a LightCycler 96 system (Roche, Basel, Switzerland). The qPCR reaction was made with 5 µl KOD SYBR (TOYOBO), 0.4 µL forward primer [10 pmol/µl], 0.4 µl reverse primer [10 pmol/µl], and 3.2 µl water. The qPCR was performed under the following conditions: 180 s at 95°C, 40 cycles of 8 s at 98°C, 10 s at 60°C, and 10 s at 68°C, followed by heating to 90°C for melting curve analysis. Mean cycle threshold (*C_T_*) values of samples were calculated for at least 5 replicates, and both Δ*C_T_* (*C_T_* Z gene – *C_T_* ef1a) and ΔΔ*C_T_* (Δ*C_T_*male – Δ*C_T_*fem) values were calculated.

### Cell-based assays

Fat bodies septically isolated from *Wolbachia*-infected *H. bolina* lines (ISG-1 and YOG-1) were placed in the cell lines AeAl2. After confirming that *Wolbachia* was stably maintained by diagnostic PCR, *Wolbachia*-positive AeAl2 cells were used as donors for *Wolbachia* transinfection into *the O. scapulalis* NARO-Ossc-M1 (OsM1) cell line. *Wolbachia*-infected AeAl2 cells were passed through a 5.0-µm filter (Cat. No. FJ25ASCCA050PL01, GVS, Via Roma, Italy), and six drops (approximately 300 µL) of the flow-through were added to the recipient OsM1 cell line.

Total RNA (200–500 ng) extracted from harvested cells using Isogen II (Nippon Gene, Tokyo, Japan) was reverse-transcribed using a PrimeScript II 1st strand cDNA Synthesis Kit (Takara Bio, Shiga, Japan) as described above. qPCR was performed as described above with primers targeting *masc*, *dsx*, and *znf2* genes (**Table S4)**. The relative expression (*Osmasc*, *Osdsx*, and *Osznf2* versus the control gene *Osef1a*) and the ratio of male-to-female splice variants were estimated.

### Transfection assays and quantification of male type *dsx* in Sf9 cells

Total RNA (1000 ng) derived from the male butterfly (*w*Bol1Y-cured YOG-1 line) was reverse-transcribed using PrimeScript™ II Reverse Transcriptase (TaKaRa, Shiga, Japan). The coding sequence of *Hbmasc* was amplified from the cDNA using KOD-one (TOYOBO) with the primer set shown in **Table S4** with following PCR conditions: 35 cycles of 98°C for 10 s, 62°C for 5 s, and 68°C for 15 s. The amplicon was purified with the QIAquick PCR purification kit (Qiagen) and cloned into the pIZ/V5-His vector using the NEB Hifi genomic assembly (New England Biolabs). We used the codon-optimized *Hb-oscar* gene, identical to the sequences of the codon optimized *Hm-oscar* (24) cloned into the pIZ/V5-His vector for the downstream assays.

To verify the masculinizing function of *Hbmasc*, Sf9 cells (4 × 10^5^ cells per dish, diameter 35 mm) were transfected with 2 µg of plasmid DNA (1 µg of pIZ/V5-His having *Hbmasc* and 1 µg of non-inserted pIZ/V5-His) using FuGENE HD (Promega, WI, USA), as described in Katsuma et al. (20). To clarify whether *Hb-oscar* suppressed the function of *Hbmasc*, 2 µg of plasmid DNA (1 µg of pIZ/V5-His bearing *Hbmasc* and 1 µg of *Hb-oscar* or non-inserted vector) was transfected to the Sf9 cells using FuGENE HD (Promega). Three days after transfection, the cells were collected and subjected to RNA extraction via TRI REAGENT® (Molecular Research Center Inc., USA) and cDNA construction with AMV transcriptase (TaKaRa). The degree of masculinization in the Sf9 cells (default female-type sex determination) was verified by quantifying the expression levels of *Sfdsx^M^*, normalized with *SfRp49* gene (**Table S4**). Co-transfection of *Hm-oscar* and *Hmmasc* was also assessed as a control. The expression levels of *Sfdsx^M^* were analyzed as described in Katsuma et al. (20). In brief, the relative expression of *Sfdsx^M^* (*C_T_ Sfdsx^M^* / *C_T_ SfRp49*) was calculated for each experimental group. Then, relative expression of *Sfdsx^M^*of each sample was normalized by setting the mean in the *Hmmasc* singly transfected group as 100. The normalized relative expression of *Sfdsx^M^* was visualized using R v4.0 with ggplot2 (https://ggplot2.tidyverse.org/).

### Statistical analysis

The sex ratio bias of adults and embryos was analyzed using the numbers of males and females with a binominal test. Hatchability (number of hatched neonates /total number of eggs) of the wBol1Y/wBol1J-infected hosts was analyzed with a Wilcoxon test in R 4.0.0.

## Data Availability

The sequence read data are publicly available in DDBJ under the accession numbers PRJDB18323 and PRJDB19678 (BioProject). *Wolbachia* genomes are available in the DDBJ database under the following accession numbers: *w*Bol1Y (SAMD00796175, AP035771) and *w*Bol1J (SAMD00796176, AP035772). Sequences for mitochondrial genome (LC850813), *Hbmasc* (LC850935), and *Hbdsx* (LC853266-LC853266) of *H. bolina* are available in the DDBJ database. Other data generated in this study are included in the manuscript and Supplementary Material.

## Supporting information

Suppremental data files

## Acknowledgments

We thank Hiromi Detani (Osaka, Japan) and Kazunari Nicho (Kagoshima, Japan) for their kind advice and suggestion. We acknowledge support from the Japan Society for the Promotion of Science (JSPS) Research Fellowships for Young Scientists [Grant Number 19J13123 and 21J00895 to H. Arai], JSPS Grant-in-Aid for Scientific Research [Grant Number 22K14902 to H. Arai, 24H02289 to S. Katsuma, and 24H02293 to D. Kageyama], JSPS Overseas Challenge Program for Young Researchers (2019) with RISTEK Foreign Research Permit [1539057329], and the Cabinet Office, Government of Japan, Moonshot Research and Development Program for Agriculture, Forestry and Fisheries (Grant Number JPJ009237).

## Author Contributions

H. Arai reared insects, conducted fieldwork, genome analysis, mating experiments, cell transinfection assays, transcriptome analysis, and data analysis; designed experiments; wrote the original manuscript; and revised the manuscript.

Arman Wijonarko organised and conducted fieldwork, reared insects, revised the manuscript and contributed to the discussion.

Susumu Katsuma conducted transfection assays using Sf9 cells, revised the manuscript and contributed to the discussion.

Hideshi Naka reared insects and helped mating experiments, revised the manuscript and contributed to the discussion.

Daisuke Kageyama revised the manuscript and contributed to the discussion. Emily A. Hornett revised the manuscript and contributed to the discussion.

Gregory D. D. Hurst revised the manuscript and contributed to the discussion.

## Inclusion and Diversity

We support inclusive, diverse, and equitable conduct of research.

## References

1. N. Gerardo, G. Hurst, Q&A: Friends (but sometimes foes) within: the complex evolutionary ecology of symbioses between host and microbes. BMC Biol. 15, 126 (2017).

2. G. D. D. Hurst, C. L. Frost, Reproductive Parasitism: Maternally Inherited Symbionts in a Biparental World. Cold Spring Harb. Perspect. Biol. 7, a017699 (2015).

3. A. E. Douglas, Nutritional Interactions in Insect-Microbial Symbioses: Aphids and Their Symbiotic Bacteria Buchnera. Annu. Rev. Entomol. 43, 17–37 (1998).

4. P. T. Hamilton, F. Peng, M. J. Boulanger, S. J. Perlman, A ribosome-inactivating protein in a Drosophila defensive symbiont. Proc. Natl. Acad. Sci. 113, 350–355 (2016).

5. K. M. Oliver, P. H. Degnan, M. S. Hunter, N. A. Moran, Bacteriophages Encode Factors Required for Protection in a Symbiotic Mutualism. Science 325, 992–994 (2009).

6. J. A. Russell, N. A. Moran, Costs and benefits of symbiont infection in aphids: variation among symbionts and across temperatures. Proc. R. Soc. B Biol. Sci. 273, 603–610 (2005).

7. C. K. Cornwallis, et al., Symbioses shape feeding niches and diversification across insects. *Nat*. Ecol. Evol. 7, 1022–1044 (2023).

8. G. D. Hurst, F. M. Jiggins, Male-killing bacteria in insects: mechanisms, incidence, and implications. Emerg. Infect. Dis. 6, 329–336 (2000).

9. L. D. Hurst, The incidences and evolution of cytoplasmic male killers. Proc. R. Soc. Lond. B Biol. Sci. 244, 91–99 (1991).

10. S. Charlat, et al., Extraordinary Flux in Sex Ratio. Science 317, 214–214 (2007).

11. M. Hayashi, M. Nomura, D. Kageyama, Rapid comeback of males: evolution of male-killer suppression in a green lacewing population. Proc. R. Soc. B Biol. Sci. 285, 20180369 (2018).

12. E. A. Hornett, et al., Evolution of Male-Killer Suppression in a Natural Population. PLOS Biol. 4, e283 (2006).

13. T. M. O. Majerus, M. E. N. Majerus, Intergenomic Arms Races: Detection of a Nuclear Rescue Gene of Male-Killing in a Ladybird. PLOS Pathog. 6, e1000987 (2010).

14. K. M. Richardson, et al., A male-killing Wolbachia endosymbiont is concealed by another endosymbiont and a nuclear suppressor. PLOS Biol. 21, e3001879 (2023).

15. K. Yoshida, S. Sanada-Morimura, S.-H. Huang, M. Tokuda, Silence of the killers: discovery of male-killing suppression in a rearing strain of the small brown planthopper, Laodelphax striatellus. Proc. R. Soc. B Biol. Sci. 288, 20202125 (2021).

16. H. Arai, et al., Prophage-encoded Hm-oscar gene recapitulates Wolbachia-induced male killing in the tea tortrix moth Homona magnanima. eLife 13 (2024).

17. H. Arai, et al., Diverse Molecular Mechanisms Underlying Microbe-Inducing Male Killing in the Moth Homona magnanima. Appl. Environ. Microbiol. 89, e02095–22 (2023).

18. T. Fukui, et al., The Endosymbiotic Bacterium Wolbachia Selectively Kills Male Hosts by Targeting the Masculinizing Gene. PLOS Pathog. 11, e1005048 (2015).

19. T. Harumoto, B. Lemaitre, Male-killing toxin in a bacterial symbiont of Drosophila. Nature 557, 252–255 (2018).

20. S. Katsuma, et al., A Wolbachia factor for male killing in lepidopteran insects. Nat. Commun. 13, 6764 (2022).

21. T. N. Sugimoto, T. Kayukawa, T. Matsuo, T. Tsuchida, Y. Ishikawa, A short, high-temperature treatment of host larvae to analyze *Wolbachia*–host interactions in the moth *Ostrinia scapulalis*. J. Insect Physiol. 81, 48–51 (2015).

22. T. N. Sugimoto, Y. Ishikawa, A male-killing Wolbachia carries a feminizing factor and is associated with degradation of the sex-determining system of its host. Biol. Lett. 8, 412–415 (2012).

23. P. M. Ferree, A. Avery, J. Azpurua, T. Wilkes, J. H. Werren, A Bacterium Targets Maternally Inherited Centrosomes to Kill Males in Nasonia. Curr. Biol. 18, 1409–1414 (2008).

24. H. Arai, et al., Combined actions of bacteriophage-encoded genes in *Wolbachia*-induced male lethality. iScience 26, 106842 (2023).

25. J. I. Perlmutter, et al., The phage gene wmk is a candidate for male killing by a bacterial endosymbiont. PLOS Pathog. 15, e1007936 (2019).

26. E. A. Hornett, D. Kageyama, G. D. D. Hurst, Sex determination systems as the interface between male-killing bacteria and their hosts. Proc. R. Soc. B Biol. Sci. 289, 20212781 (2022).

27. M. A. Brockhurst, et al., Running with the Red Queen: the role of biotic conflicts in evolution. Proc. R. Soc. B Biol. Sci. 281, 20141382 (2014).

28. E. A. Dyson, M. K. Kamath, G. D. D. Hurst, Wolbachia infection associated with all-female broods in Hypolimnas bolina (Lepidoptera: Nymphalidae): evidence for horizontal transmission of a butterfly male killer. Heredity 88, 166–171 (2002).

29. E. A. Dyson, G. D. D. Hurst, Persistence of an extreme sex-ratio bias in a natural population. Proc. Natl. Acad. Sci. 101, 6520–6523 (2004).

30. E. B. Poulton, All female families of Hypolimnas bolina, bred in Fiji by HW Simmonds. Proc R Entomol Soc Lond 1923, 9–12 (1923).

31. G. H. E. Hopkins, Insects of Samoa and Other Samoan Terrestrial Arthropoda (1927).

32. W. Mitsuhashi, H. Fukuda, K. Nicho, R. Murakami, Male-killing Wolbachia in the butterfly Hypolimnas bolina. Entomol. Exp. Appl. 112, 57–64 (2004).

33. C. Clarke, P. M. Sheppard, V. Scali, All-female broods in the butterfly Hypolimnas bolina (L.). Proc. R. Soc. Lond. B Biol. Sci. 189, 29–37 (1975).

34. H. Fukuda, K. Nicho, Some Problems in the Great Egg-fly, Hypolimnas bolina L. (Nymphalidae). Spec. Bull. Lepidopterol. Soc. Jpn. 6, 35–68 (1988).

35. H. Ikeda, Memories on Hypolimnas bolina (3) notes on the sex-ratio. Yadoriga 166, 29–36 (1996).

36. W. Mitsuhashi, H. Ikeda, M. Muraji, Fifty-year trend towards suppression of Wolbachia-induced male-killing by its butterfly host, Hypolimnas bolina. J. Insect Sci. 11, 92 (2011).

37. S. Charlat, et al., The joint evolutionary histories of Wolbachia and mitochondria in Hypolimnas bolina. BMC Evol. Biol. 9, 64 (2009).

38. E. A. Hornett, S. Charlat, N. Wedell, C. D. Jiggins, G. D. D. Hurst, Rapidly Shifting Sex Ratio across a Species Range. Curr. Biol. 19, 1628–1631 (2009).

39. F. M. Jiggins, G. D. D. Hurst, M. E. N. Majerus, Sex-ratio-distorting Wolbachia causes sex-role reversal in its butterfly host. Proc. R. Soc. Lond. B Biol. Sci. 267, 69–73 (2000).

40. S. Charlat, et al., Competing Selfish Genetic Elements in the Butterfly Hypolimnas bolina. Curr. Biol. 16, 2453–2458 (2006).

41. M. F. Richardson, et al., Population Genomics of the Wolbachia Endosymbiont in Drosophila melanogaster. PLOS Genet. 8, e1003129 (2012).

42. M. Scholz, et al., Large scale genome reconstructions illuminate Wolbachia evolution. Nat. Commun. 11, 5235 (2020).

43. A. Duplouy, et al., Draft genome sequence of the male-killing Wolbachia strain wBol1 reveals recent horizontal gene transfers from diverse sources. BMC Genomics 14, 20 (2013).

44. A. Orteu, et al., Optix and cortex/ivory/mir-193 again: the repeated use of two mimicry hotspot loci. Proc. R. Soc. B Biol. Sci. 291, 20240627 (2024).

45. E. A. Hornett, et al., You can’t keep a good parasite down: evolution of a male-killer suppressor uncovers cytoplasmic incompatibility. Evolution 62, 1258–1263 (2008).

46. M. G. Suzuki, “Sex Determination Cascade in Insects: A Great Treasure House of Alternative Splicing” in Reproductive and Developmental Strategies: The Continuity of Life, K. Kobayashi, T. Kitano, Y. Iwao, M. Kondo, Eds. (Springer Japan, 2018), pp. 267–288.

47. T. Kiuchi, et al., A single female-specific piRNA is the primary determiner of sex in the silkworm. Nature 509, 633–636 (2014).

48. G. Gopinath, K. P. Arunkumar, K. Mita, J. Nagaraju, Role of *Bmznf-2*, a *Bombyx mori* CCCH zinc finger gene, in masculinisation and differential splicing of *Bmtra-*2. Insect Biochem. Mol. Biol. 75, 32–44 (2016).

49. H. Arai, et al., Cell-based assays and comparative genomics revealed the conserved and hidden effects of Wolbachia on insect sex determination. PNAS Nexus 3, pgae348 (2024).

50. B. Herran, et al., Cell-based analysis reveals that sex-determining gene signals in Ostrinia are pivotally changed by male-killing Wolbachia. PNAS Nexus 2, pgac293 (2023).

51. J. L. Vaughn, R. H. Goodwin, G. J. Tompkins, P. McCawley, The establishment of two cell lines from the insect Spodoptera frugiperda (Lepidoptera; Noctuidae). In Vitro 13, 213–217 (1977).

52. E. A. Hornett, et al., The Evolution of Sex Ratio Distorter Suppression Affects a 25 cM Genomic Region in the Butterfly Hypolimnas bolina. PLOS Genet. 10, e1004822 (2014).

53. L. A. Reynolds, E. A. Hornett, C. D. Jiggins, G. D. D. Hurst, Suppression of Wolbachia-mediated male-killing in the butterfly Hypolimnas bolina involves a single genomic region. PeerJ 7, e7677 (2019).

54. H. Arai, M. N. Inoue, D. Kageyama, Male-killing mechanisms vary between Spiroplasma species. Front. Microbiol. 13 (2022).

55. D. Kageyama, et al., A male-killing gene encoded by a symbiotic virus of Drosophila. Nat. Commun. 14, 1357 (2023).

56. K. Nagamine, et al., Male-killing virus in a noctuid moth Spodoptera litura. Proc. Natl. Acad. Sci. 120, e2312124120 (2023).

57. H. Arai, M. Watada, D. Kageyama, Two male-killing Wolbachia from Drosophila birauraia that are closely related but distinct in genome structure. R. Soc. Open Sci. 11, 231502 (2024).

58. G. D. D. Hurst, et al., Male–killing Wolbachia in two species of insect. Proc. R. Soc. Lond. B Biol. Sci. 266, 735–740 (1999).

59. W. D. Hamilton, Extraordinary Sex Ratios: A sex-ratio theory for sex linkage and inbreeding has new implications in cytogenetics and entomology. Science 156, 477– 488 (1967).

60. J. D. G. Jones, J. L. Dangl, The plant immune system. Nature 444, 323–329 (2006).

61. C. Courret, C.-H. Chang, K. H.-C. Wei, C. Montchamp-Moreau, A. M. Larracuente, Meiotic drive mechanisms: lessons from Drosophila. Proc. R. Soc. B Biol. Sci. 286, 20191430 (2019).

62. S. A. Frank, The Evolutionary Dynamics of Cytoplasmic Male Sterility. Am. Nat. 133, 345–376 (1989).

63. D. E. Perez, C. I. Wu, Further characterization of the Odysseus locus of hybrid sterility in Drosophila: one gene is not enough. Genetics 140, 201–206 (1995).

64. C.-T. Ting, S.-C. Tsaur, M.-L. Wu, C.-I. Wu, A Rapidly Evolving Homeobox at the Site of a Hybrid Sterility Gene. Science 282, 1501–1504 (1998).

65. H. Arai, et al., Conserved infections and reproductive phenotypes of Wolbachia symbionts in Asian tortrix moths. Environ. Microbiol. Rep. 16, e13219 (2024).

66. C. Clarke, P. M. Sheppard, The genetics of the mimetic butterfly Hypolimnas bolina (L.). Philos. Trans. R. Soc. Lond. B Biol. Sci. 272, 229–265 (1975).

67. H. Arai, et al., Multiple Infection and Reproductive Manipulations of Wolbachia in Homona magnanima (Lepidoptera: Tortricidae). Microb. Ecol. 77, 257–266 (2019).

68. T. Magoč, S. L. Salzberg, FLASH: fast length adjustment of short reads to improve genome assemblies. Bioinformatics 27, 2957–2963 (2011).

69. E. Bolyen, et al., Reproducible, interactive, scalable and extensible microbiome data science using QIIME 2. Nat. Biotechnol. 37, 852–857 (2019).

70. C. Quast, et al., The SILVA ribosomal RNA gene database project: improved data processing and web-based tools. Nucleic Acids Res. 41, D590–596 (2013).

71. M. Kolmogorov, J. Yuan, Y. Lin, P. A. Pevzner, Assembly of long, error-prone reads using repeat graphs. Nat. Biotechnol. 37, 540–546 (2019).

72. B. J. Walker, et al., Pilon: An Integrated Tool for Comprehensive Microbial Variant Detection and Genome Assembly Improvement. PLOS ONE 9, e112963 (2014).

73. H. Li, Minimap2: pairwise alignment for nucleotide sequences. Bioinformatics 34, 3094–3100 (2018).

74. Y. Tanizawa, T. Fujisawa, Y. Nakamura, DFAST: a flexible prokaryotic genome annotation pipeline for faster genome publication. Bioinformatics 34, 1037–1039 (2018).

75. M. J. Sullivan, N. K. Petty, S. A. Beatson, Easyfig: a genome comparison visualizer. Bioinformatics 27, 1009–1010 (2011).

76. C. Chen, et al., TBtools: An Integrative Toolkit Developed for Interactive Analyses of Big Biological Data. Mol. Plant 13, 1194–1202 (2020).

77. J. Martinez, L. Klasson, J. J. Welch, F. M. Jiggins, Life and Death of Selfish Genes: Comparative Genomics Reveals the Dynamic Evolution of Cytoplasmic Incompatibility. Mol. Biol. Evol. 38, 2–15 (2021).

78. H. Li, et al., The Sequence Alignment/Map format and SAMtools. Bioinformatics 25, 2078–2079 (2009).

79. M. A. Larkin, et al., Clustal W and Clustal X version 2.0. Bioinformatics 23, 2947– 2948 (2007).

80. L. Baldo, et al., Multilocus Sequence Typing System for the Endosymbiont Wolbachia pipientis. Appl. Environ. Microbiol. 72, 7098–7110 (2006).

81. D. M. Emms, S. Kelly, OrthoFinder: phylogenetic orthology inference for comparative genomics. Genome Biol. 20, 238 (2019).

82. W. Shen, S. Le, Y. Li, F. Hu, SeqKit: A Cross-Platform and Ultrafast Toolkit for FASTA/Q File Manipulation. PLOS ONE 11, e0163962 (2016).

83. K. Katoh, K. Kuma, H. Toh, T. Miyata, MAFFT version 5: improvement in accuracy of multiple sequence alignment. Nucleic Acids Res. 33, 511–518 (2005).

84. S. Capella-Gutiérrez, J. M. Silla-Martínez, T. Gabaldón, trimAl: a tool for automated alignment trimming in large-scale phylogenetic analyses. Bioinformatics 25, 1972– 1973 (2009).

85. J. Rozas, et al., DnaSP 6: DNA Sequence Polymorphism Analysis of Large Data Sets. Mol. Biol. Evol. 34, 3299–3302 (2017).

86. H. J. Bandelt, P. Forster, A. Röhl, Median-joining networks for inferring intraspecific phylogenies. Mol. Biol. Evol. 16, 37–48 (1999).

87. A. M. Bolger, M. Lohse, B. Usadel, Trimmomatic: a flexible trimmer for Illumina sequence data. Bioinformatics 30, 2114–2120 (2014).

88. M. G. Grabherr, et al., Trinity: reconstructing a full-length transcriptome without a genome from RNA-Seq data. Nat. Biotechnol. 29, 644–652 (2011).

89. N. L. Bray, H. Pimentel, P. Melsted, L. Pachter, Near-optimal probabilistic RNA-seq quantification. Nat. Biotechnol. 34, 525–527 (2016).

