## Supplementary material for "Evolution of *Wolbachia* male-killing mechanism within a host species": Suppremental data files

**Supplemental Figures and Tables**

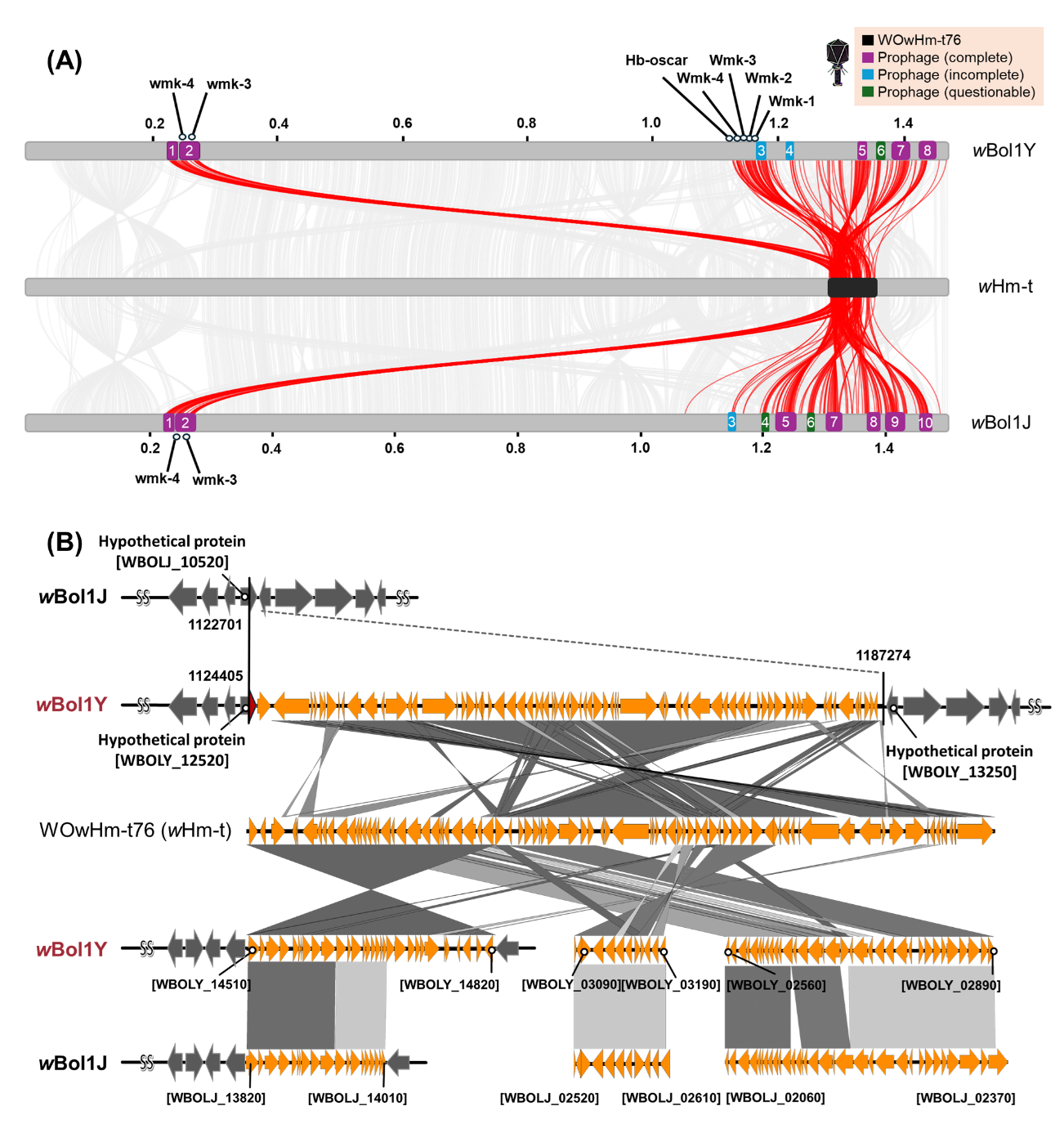

**Fig.S1 Prophage regions in *w*Bol1Y and *w*Bol1J and their homology to WO*w*Hm-t76**

(A) Prophage insertions in the *w*Bol1J and *w*Bol1Y genomes. Prophage regions were annotated using PHASTEST, and the region that showed homologies with the WO*w*Hm-t76 region (show with black) is connected and highlighted with a red line. Numbers on/under the genome indicate the position of the genome (e.g. 0.2 Mb). (B) Homologies between the WO*w*Hm-t76 and *w*Bol1 genomes. In addition to the *w*Bol1Y-specific insertion (ca. 63 kb, with high homology to 23-76 kb of WO*w*Hm-t76), a prophage region (ca. 23 kb) with high homology to the 1-23 kb of WO*w*Hm-t76 was identified in the *w*Bol1Y genome. *w*Bol1J and *w*Bol1Y shared several prophage regions with high homology to WO*w*Hm-t76.

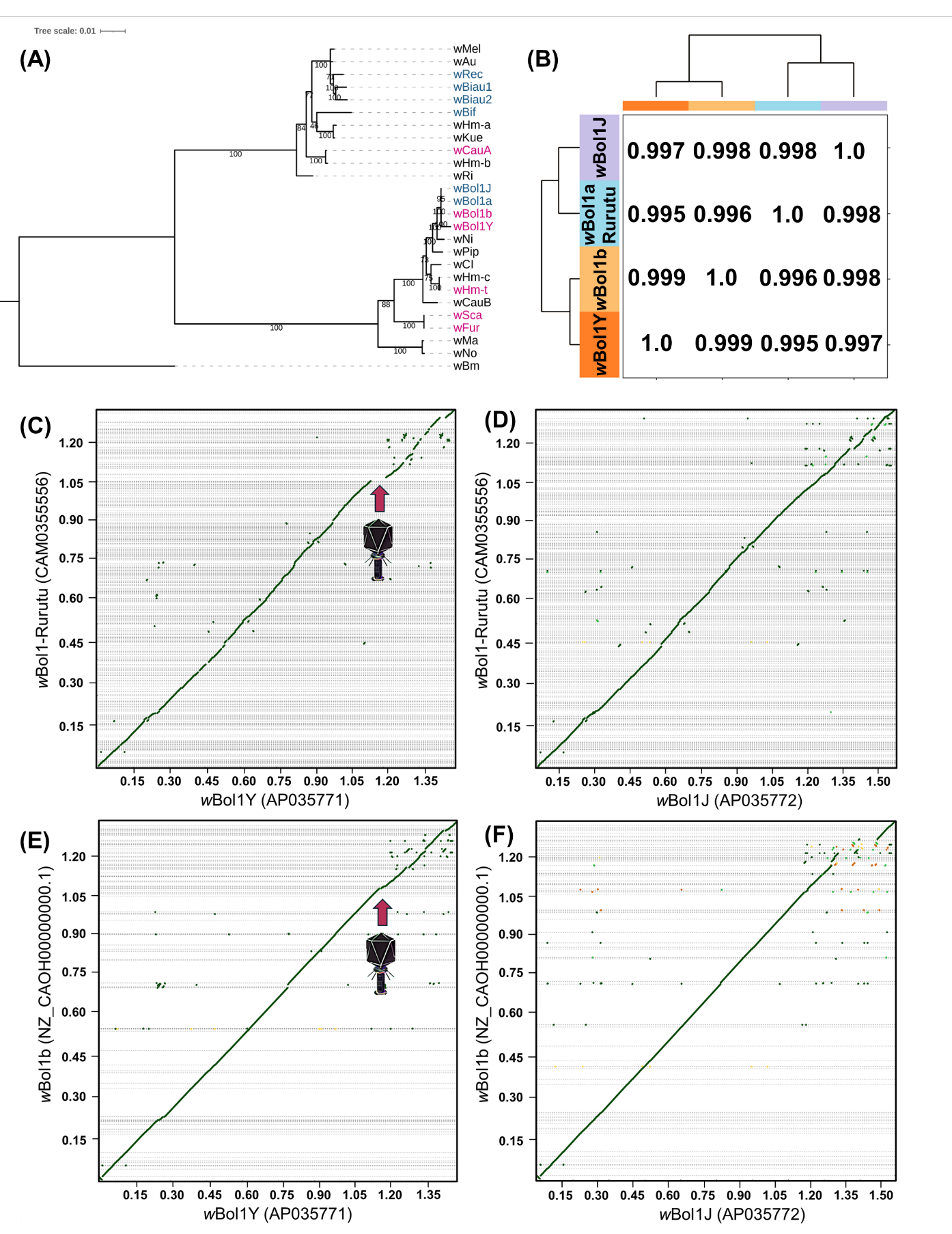

**Fig. S2. Previously described MK *w*Bol1 strain lacked the *Hb-oscar*-bearing prophage element and shared a more recent origin with the suppressed *w*Bol1J.**
(A) Phylogenetic tree based on single-copy *Wolbachia* genes. MK *Wolbachia* strains harboring *oscar* homologs are highlighted in magenta, and *oscar*-deficient strains are highlighted in blue. (B) ANI values and phylogenetic relationships of *w*Bol1 strains in *H. bolina*. (C-F) Harr plots of the genomes of *w*Bol1 (MK, Rurutu), *w*Bol1b (Moorea), *w*Bol1Y (MK, Yogyakarta), and *w*Bol1J (suppressed MK, Ishigaki). The homology of their genomes is highlighted in green (>80% identity), yellow (50-80% identity), and orange (30-50% identity). The fragmented genome of *w*Bol1 (MK, Rurutu) and *w*Bol1b (Moorea) were aligned against the reference genomes of *w*Bol1Y (C and E) and *w*Bol1J (D and F). The *w*Bol1Y-specific prophage insertion carrying the *Hb-oscar* gene is indicated by an arrow with a phage illustration. Numbers along the genome represent genomic positions (e.g., 0.2 Mb).

**Table S1 Bacterial communities of the *w*Bol1-infected host lines**

| Family | Genus | Bacterial abundance (read counts) | |
| --- | --- | --- | --- |
|  |  | ISG (non-MK) | YOG (MK) |
| Anaplasmataceae | *Wolbachia* | 24731 | 27844 |
| Acetobacteraceae | *Asaia* | 855 | 2012 |
| Weeksellaceae | *Apibacter* | 4309 | 1441 |
| Yersiniaceae | *Serratia* | 0 | 82 |
| Propionibacteriaceae | *Cutibacterium* | 0 | 27 |
| Sphingomonadaceae | *Sphingomonas* | 27 | 22 |
| Orbaceae | *Orbus* | 1938 | 17 |
| Corynebacteriaceae | *Lawsonella* | 14 | 12 |
| Dermacoccaceae | *Dermacoccus* | 0 | 12 |
| Acetobacteraceae | *Roseomonas* | 0 | 7 |
| Lactobacillaceae | *Lactobacillus* | 97 | 0 |
| Rhizobiaceae | *Bartonella* | 2397 | 0 |
| Acetobacteraceae | *Commensalibacter* | 191 | 0 |
| Dysgonomonadaceae | *Dysgonomonas* | 1550 | 0 |
| Enterobacteriaceae | Unclassified | 52 | 0 |
| Enterococcaceae | *Enterococcus* | 38 | 0 |
| Staphylococcaceae | *Staphylococcus* | 20 | 0 |
| Alcaligenaceae | *Achromobacter* | 8 | 0 |
| Pseudomonadaceae | *Pseudomonas* | 5 | 0 |

**Table S2 Phenotype-associated *Wolbachia* genes and homologies based on BLASTn searches.**

| Query | *w*Bol1 genes | Identity | Query | | *Wolbachia* protein | | E-value | Bit score |
| --- | --- | --- | --- | --- | --- | --- | --- | --- |
|  |  |  | Start | End | Start | End |  |  |
| wHm-t [Hb-oscar] | WBOLY_12540 | 100 | 1 | 3546 | 1 | 3546 | 0 | 6549 |
| wBol1_pair1 [Type_I cifA] | WBOLJ_13580 | 100 | 1 | 1476 | 1 | 1476 | 0 | 2726 |
| wBol1_pair2 [Type_IV cifA] | WBOLJ_11740 | 100 | 1 | 1338 | 1 | 1338 | 0 | 2471 |
| wBol1_pair1 [Type_I cifA] | WBOLY_14240 | 99.932 | 1 | 1476 | 1 | 1475 | 0 | 2719 |
| wBol1_pair2 [Type_IV cifA] | WBOLY_15570 | 100 | 1 | 1338 | 1 | 1338 | 0 | 2471 |
| wBol1_pair2 [Type_IV cifA] | WBOLY_14040 | 100 | 1 | 1338 | 1 | 1338 | 0 | 2471 |
| wBol1_pair1 [Type_I cifB] | WBOLJ_13570 | 100 | 1 | 3525 | 1 | 3525 | 0 | 6510 |
| wBol1_pair2 [Type_IV cifB] | WBOLJ_11730 | 100 | 1 | 2199 | 1 | 2199 | 0 | 4061 |
| wBol1_pair1 [Type_I cifB] | WBOLY_14230 | 100 | 1 | 3525 | 1 | 3525 | 0 | 6510 |
| wBol1_pair2 [Type_IV cifB] | WBOLY_15560 | 99.896 | 1 | 1916 | 1 | 1914 | 0 | 3526 |
| wBol1_pair2 [Type_IV cifB] | WBOLY_14030 | 99.896 | 1 | 1916 | 1 | 1914 | 0 | 3526 |
| wHm-t [wmk4] | WBOLJ_02550 | 100 | 1 | 1005 | 1 | 1005 | 0 | 1857 |
| wHm-t [wmk3] | WBOLJ_02560 | 100 | 1 | 897 | 1 | 897 | 0 | 1657 |
| wMel WD0626 | WBOLJ_02610 | 73.235 | 64 | 912 | 82 | 939 | 2.77E-74 | 274 |
| wHm-t [wmk3] | WBOLJ_11130 | 84.32 | 31 | 360 | 604 | 939 | 3.73E-89 | 322 |
| wHm-t [wmk3] | WBOLJ_11160 | 86.942 | 1 | 885 | 13 | 903 | 0 | 992 |
| wHm-t [wmk4] | WBOLJ_11170 | 93.098 | 42 | 692 | 33 | 683 | 0 | 953 |
| wMel WD0626 | WBOLJ_11800 | 85.965 | 1 | 569 | 1 | 561 | 1.12E-172 | 601 |
| wHm-t [wmk3] | WBOLJ_11810 | 98.75 | 28 | 187 | 4 | 163 | 4.89E-78 | 285 |
| wHm-t [wmk2] | WBOLJ_11820 | 91.667 | 217 | 360 | 4 | 138 | 1.10E-49 | 191 |
| wHm-t [wmk3] | WBOLJ_11880 | 88.787 | 1 | 864 | 13 | 885 | 0 | 1061 |
| wHm-t [wmk4] | WBOLJ_11890 | 89.466 | 42 | 692 | 33 | 683 | 0 | 821 |
| wMel WD0626 | WBOLJ_12800 | 85.246 | 64 | 912 | 73 | 921 | 0 | 870 |
| wHm-t [wmk3] | WBOLJ_12860 | 88.826 | 1 | 879 | 1 | 882 | 0 | 1077 |
| wHm-t [wmk4] | WBOLJ_12870 | 94.239 | 34 | 709 | 25 | 699 | 0 | 1031 |
| wMel WD0626 | WBOLJ_13600 | 87.991 | 1 | 912 | 1 | 912 | 0 | 1075 |
| wHm-t [wmk3] | WBOLJ_13630 | 88.939 | 1 | 879 | 1 | 882 | 0 | 1083 |
| wHm-t [wmk4] | WBOLJ_13640 | 92.121 | 34 | 692 | 25 | 683 | 0 | 929 |
| wHm-t [wmk4] | WBOLY_03120 | 100 | 1 | 1005 | 1 | 1005 | 0 | 1857 |
| wHm-t [wmk3] | WBOLY_03130 | 100 | 1 | 897 | 1 | 897 | 0 | 1657 |
| wHm-t [wmk3] | WBOLY_03190 | 80.119 | 31 | 360 | 211 | 546 | 7.86E-66 | 244 |
| wHm-t [wmk4] | WBOLY_12770 | 100 | 1 | 1005 | 1 | 1005 | 0 | 1857 |
| wHm-t [wmk3] | WBOLY_12780 | 100 | 1 | 897 | 1 | 897 | 0 | 1657 |
| wHm-t [wmk2] | WBOLY_12890 | 100 | 1 | 360 | 1 | 360 | 0 | 665 |
| wHm-t [wmk1] | WBOLY_12900 | 100 | 1 | 561 | 1 | 561 | 0 | 1037 |
| wMel WD0626 | WBOLY_14260 | 87.882 | 1 | 912 | 1 | 912 | 0 | 1070 |
| wHm-t [wmk3] | WBOLY_14290 | 88.939 | 1 | 879 | 1 | 882 | 0 | 1083 |
| wHm-t [wmk4] | WBOLY_14300 | 92.121 | 34 | 692 | 25 | 683 | 0 | 929 |
| wHm-t [wmk3] | WBOLY_14880 | 84.32 | 31 | 360 | 172 | 507 | 3.53E-89 | 322 |
| wHm-t [wmk3] | WBOLY_14920 | 86.942 | 1 | 885 | 13 | 903 | 0 | 992 |
| wHm-t [wmk4] | WBOLY_14930 | 93.098 | 42 | 692 | 33 | 683 | 0 | 953 |

**Table S3 Statistical data for the quantification of Z-linked gene expression**

| Comparisons | T-value | P-value |
| --- | --- | --- |
| Free-F vs Free-M | 0.394 | 0.998 |
| Free-F vs wBol1J-F | 0.402 | 0.998 |
| Free-F vs wBol1J-M | 1.267 | 0.802 |
| Free-F vs wBol1Y-F | 0.032 | 1.000 |
| Free-F vs wBol1Y-M | 3.942 | 0.001 |
| Free-M vs wBol1J-F | 0.804 | 0.966 |
| Free-M vs wBol1J-M | 1.478 | 0.677 |
| Free-M vs wBol1Y-F | 1.215 | 0.829 |
| Free-M vs wBol1Y-M | 3.796 | 0.002 |
| wBol1J-F vs wBol1J-M | 0.596 | 0.991 |
| wBol1J-F vs wBol1Y-F | 0.100 | 0.999 |
| wBol1J-F vs wBol1Y-M | 3.423 | 0.008 |
| wBol1J-M vs wBol1Y-F | 0.493 | 0.996 |
| wBol1J-M vs wBol1Y-M | 4.033 | 0.001 |
| wBol1Y-F vs wBol1Y-M | 4.029 | 0.001 |

**Table S4 Primers used in this study**

| Target | Primers | Primer sequences (5'-3') |
| --- | --- | --- |
| *OsDsxM* | OsDsxMqF1 | GAGGAAGATTGATGAAGCCCAC |
|  | OsDsxMqR1 | GACGGAGGCTCTGATGACTC |
| *OsDsxF* | OsDsxFqF3 | CGAGGAAGATTGATGAAGGGAAG |
|  | OsDsxFqR2 | GCTTTGCAGCATTTTCTGGC |
| *OsMascM* | OsMascMqF1 | GCCAAATGGACATTACAACCAGTAC |
|  | OsMascMqR1 | CACGACTCGTGTCGACCAAAC |
| *OsMascF* | OsMascFqF1b | TTATATCAGGGGTGGCCTACT |
|  | OsMascFqR3 | TCAACTATATAATTCAGGTGTGGC |
| *OsZnf2M* | OsZnf2MqF1 | CCACCGAATCAAGCAATTGC |
|  | OsZnf2MqR1 | TTTTCGTTCGCTTCGGATATTTG |
| *OsZnf2F* | OsZnf2FqF3 | AGTGTTCTGTGGTAATTAATTCGC |
|  | OsZnf2FqR2 | GGGCGCGGCCGTTGATTC |
| *OsEf1a* | OsEF1a F | GACTCCGGCAAGTCCACCAC |
|  | OsEF1a R | CCTGGGCCTCCTTCTCGAAT |
| *Hb-oscar* | WHMT_00358_f | ATGATTGAAGATAGAAATGTTCCTTTATCC |
|  | WHMT_00358_r | CTACCTACCGCCTTTACCTTTGCTA |
|  | Hm/Hb/Eh-Oscar.qF | GGACCAGTAGAAGCACCTATA |
|  | Hm/Hb/Eh-Oscar.qR | AACACGACTAGCTTCAGGAC |
|  | Hm/Hb/Eh-probe | 5'6-FAM™/TTCTAAACT/ZEN™/CGACTGTGGCAGGCC/3'IB®FQP |
| *wsp* | wspF81 | TGGTCCAATAAGTGATGAAGAAAC |
|  | wspR691 | TGGAGTAGCGTTTAATTTTT |
| *gatB* | gatB_F1 | GAKTTAAAYCGYGCAGGBGTT |
|  | gatB_R1 | TGGYAAYTCRGGYAAAGATGA |
| *coxA* | coxA_F1 | TTGGRGCRATYAACTTTATAG |
|  | coxA_R1 | CTAAAGACTTTKACRCCAGT |
| *hcpA* | hcpA_F1 | GAAATARCAGTTGCTGCAAA |
|  | hcpA_R1 | GAAAGTYRAGCAAGYTCTG |
| *ftsZ* | ftsZ_F1 | ATYATGGARCATATAAARGATAG |
|  | ftsZ_R1 | TCRAGYAATGGATTRGATAT |
| *fbpA* | fbpA_F1 | GCTGCTCCRCTTGGYWTGAT |
|  | fbpA_R1 | CCRCCAGARAAAAYYACTATTC |
| *COI* | LepF: | ATTCAACCAATCATAAAGATATTGG |
|  | LepR: | TAAACTTCTGGATGTCCAAAAAATCA |
| *Hbmasc* | piZ.Hind3.F | tctgttcgaatttaaagcttCAACATGAACAATAAAAATGAACAAAATG |
|  | pIZ.BamH1.R | cacactggactagtggatccTTACTGGCGGCGCTGGTAGTAGGGC |
| *Hbdsx* | Hbdsx.Ex1.503-522F1 | CGCCTTGGAGTCTGGTATCC |
|  | Hbdsx.Ex7.40-60R1 | TTGATGTCGTCGATGGGGAC |
|  | HbExon3.123F | GCGTCGCGGAAGATAGATGA |
|  | HbExon7.51-70R | TGCCACTCCTTGATGTCGTC |
| *Hbkettin* | HbKettin-qF2.525 | CAACTTCGGCTACGTCTCCC |
|  | HbKettin-qR2.690 | ACGTTGTTCCGGTATGCCAA |
| *Hbef1a* | HbEf1a-qF2.1046 | ACCCCGGTCAAATCTCCAAC |
|  | HbEf1a-qR2.1205 | ACAATGGCAGCATCACCAGA |
| *Sfdsx^M^* | Sfdsx_qF1M | GGAAAATAGACGAAGCCCAC |
|  | Sfdsx_qR2 | CGTACTCCGTGAAGCACATG |
| SfRp49 | Sfrp49_qF1 | CCCAACATTGGTTACGGATC |
|  | Sfrp49_qR1 | TTCTTTGAGGAGACTCCGTG |
